# Multimodal identification of the mouse brain using simultaneous Ca^2+^ imaging and fMRI

**DOI:** 10.1101/2024.05.24.594620

**Authors:** Francesca Mandino, Corey Horien, Xilin Shen, Gabriel Desrosiers-Grégoire, Wendy Luo, Marija Markicevic, R. Todd Constable, Xenophon Papademetris, Mallar M. Chakravarty, Richard F. Betzel, Evelyn M.R. Lake

## Abstract

Individual differences in neuroimaging are of interest to clinical and cognitive neuroscientists based on their potential for guiding the personalized treatment of various heterogeneous neurological conditions and diseases. Despite many advantages, the workhorse in this arena, BOLD (blood-oxygen-level-dependent) functional magnetic resonance imaging (fMRI) suffers from low spatiotemporal resolution and specificity as well as a propensity for noise and spurious signal corruption. To better understand individual differences in BOLD-fMRI data, we can use animal models where fMRI, alongside complementary but more invasive contrasts, can be accessed. Here, we apply simultaneous wide-field fluorescence calcium imaging and BOLD-fMRI in mice to interrogate individual differences using a connectome-based identification framework adopted from the human fMRI literature. This approach yields high spatiotemporal resolution cell-type specific signals (here, from glia, excitatory, as well as inhibitory interneurons) from the whole cortex. We found mouse multimodal connectome-based identification to be successful and explored various features of these data.

## 1. INTRODUCTION

Individual differences are of growing interest in clinical and cognitive neuroscience^1,2^, joining more established examples of precision medicine, including personalized treatments in oncology^3^ and cardiovascular disease^4^. For nearly a decade, we have known that functional connectivity (FC) by means of non-invasive BOLD (blood-oxygen-level-dependent) functional magnetic resonance imaging (fMRI) exhibits individual-specific features that make a person’s brain distinguishable from others^5^. This framework identifies subjects from a pool using their connectome (a summary of inter-regional BOLD signal (a)synchronies or connectivity strengths). High rates of correct identification (ID) are typical in human populations (e.g., 88% in **Fig. 1**)^5–17^.

**Fig. 1.**
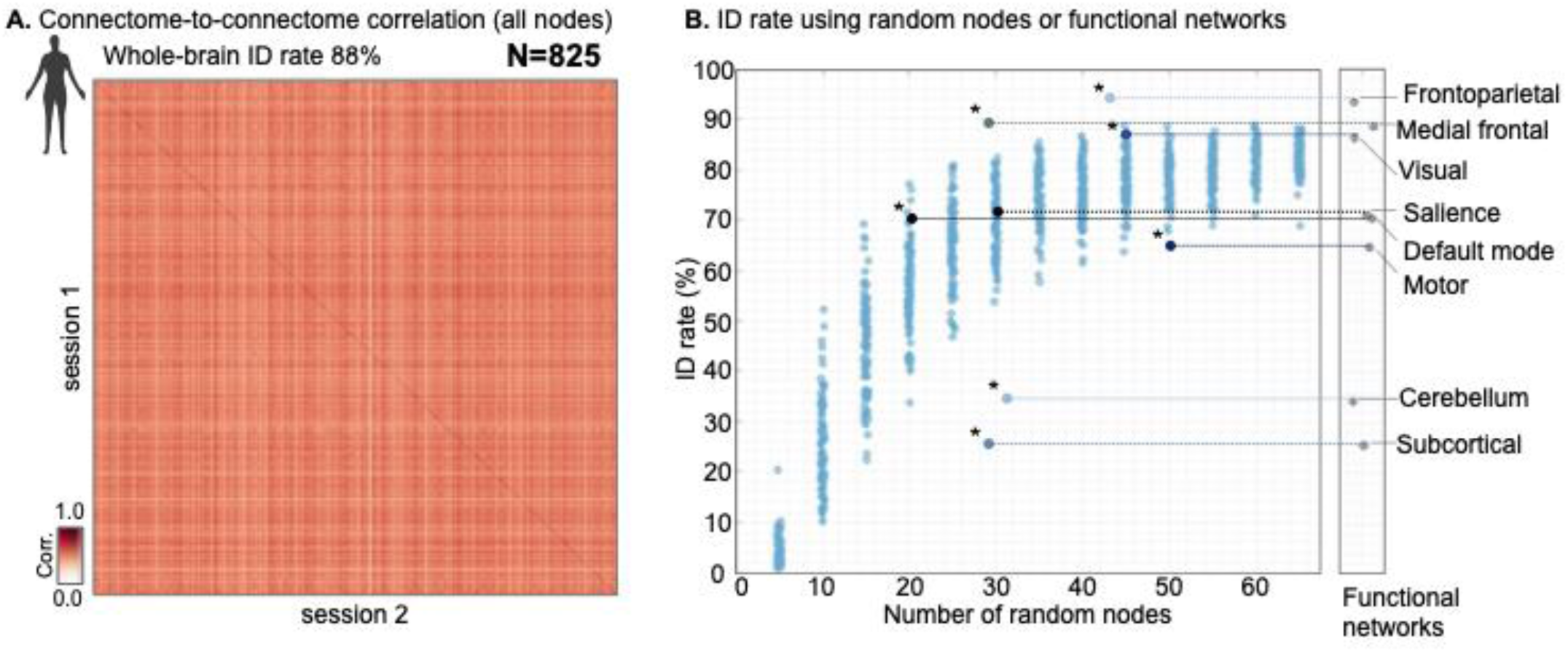
Connectome ID framework using random nodes and functional networks in humans. Connectomes were computed for human subjects from the Human Connectome Project^18^ using the Shen-268 atlas following established methods^5,19^(see ‘Human BOLD-fMRI data’ in Methods**). A.** Connectome-to-connectome correlation (Pearson) across imaging sessions. The clear diagonal indicates that connectomes from the same individual (collected during different sessions) are more similar (have a higher correlation value) than data from unmatched individuals (in-line with previous work). **B.** The Shen-268^5^ atlas has been sub-divided into 8 functional networks: (1) frontoparietal, (2) medial frontal, (3) visual, (4) salience, (5) subcortical, (6) cerebellum, (7) motor, and (8) default mode. These networks contain 20-50 nodes each. We repeated a ‘winner-take-all’ ID procedure using each functional network, and random selections of ‘x’ nodes (N = 100 iterations). ‘x’ was varied between 5 & 65 in increments of 5. As the number of randomly selected nodes increased, the ID rate also increased (reaching a plateau by approximately 40 nodes, or only 15% of all nodes). Relative to collections of randomly selected nodes, functional networks showed variable ID rates. Some performed worse (e.g., the subcortical and cerebellum networks), while others performed better (e.g., medial frontal and frontoparietal networks) than expected based on their size. Thus, as has been shown previously^5,13^, we affirmed that ID rates using BOLD-fMRI data from humans required only a small fraction of the whole-brain and showed clear functional network dependence.

This discovery garnered widespread attention by spawning the notion that clinically accessible measures of brain connectivity could be used to guide the personalized care of various neurological conditions and diseases^20^. Yet, to-date, there are no applications within the clinic. In part, this lack of translation is due to our limited understanding of the biological basis for BOLD-fMRI measures of brain connectivity, which stem from the BOLD signal being a relatively low-spatiotemporal resolution summary metric of brain functional synchrony (or lack thereof) accessed via neurogliovascular coupling. These limitations, and the lack of cell-type specificity afforded by the BOLD signal, make it clear that additional imaging modalities are needed.

To address knowledge gaps, BOLD-fMRI applied in animal models allows for invasive techniques (not applicable in humans^21^) to offer complementary measures of brain activity, and the opportunity for manipulations that can help to determine causal relationships^22,23^. Despite this potential (and some promising leads^24,25^), animal studies investigating individual differences are rare. Instead, animal studies have typically focused on group averages, which leverages the tight control animal work has over environment and genetic variability but fails to capture the clear heterogeneity that is evident in the human conditions being modeled^26–28^. Encouragingly, recent pioneering work by Bergmann et al. in 2020 established that laboratory mice, despite being homogenized by design, are identifiable using the same BOLD-fMRI connectome-based ID framework as has been used on human BOLD-fMRI data^5,29^. To the best of our knowledge, this finding has not been replicated or investigated further, until the present work.

We use a unique simultaneous wide-field calcium (WF-Ca^2+^) and BOLD-fMRI acquisition system^30^ to investigate connectome-based ID in mice using multimodal neuroimaging data. To gain insight into relationships between the BOLD signal and the underlying cellular activity, the use of optical modes, which report on brain activity using genetically encoded or virally transfected fluorescent indicators (e.g., GCaMP), yields a complementary high specificity and spatiotemporal resolution signal source^31,32^. In particular, the ability to distinguish between different neuronal and glial activities which can provide insights that fMRI (in isolation) cannot. Thus, by combining two complementary modalities, we can validate and enhance the interpretability of fMRI-based connectivity findings. Here, we implement simultaneous WF-Ca^2+^ and BOLD-fMRI in healthy adult transgenic mice (N = 45) expressing GCaMP in one of five cell types (herein called ‘groups’): (1) excitatory neurons (’SLC’), (2) glia, as well as inhibitory interneurons expressing (3) parvalbumin (‘PV’), (4) somatostatin (‘SOM’), or (5) vasoactive intestinal (poly)peptide (‘VIP’). Mice were imaged three times (∼7-days apart) under low-dose isoflurane anesthesia. Please note, that the effects of anesthesia on our findings are discussed in detail below. The overarching objective of this work was to explore connectome-based ID in mice using these unique multimodal data and to uncover the first evidence of any inter-modality convergence.

This is the first study to apply a connectome based-ID framework using simultaneously obtained multimodal data. That we found significant rates of correct IDs in both modalities affirms that connectome-based ID is likely driven by cellular activity, rather than noise. Given the markedly differing rates of ID across canonical networks in human BOLD-fMRI studies (**Fig. 1**)^5,8,12^, we investigate whether canonical networks play similar roles in mouse data as well as an edge-based approach^29,33^. The group structure of our dataset (based on cell-type expressing GCaMP) allowed us to investigate group membership, in addition to individual mouse ID rates. Data from both modalities resulted in significant group IDs – with an expected discrepancy in performance (WF-Ca^2+^ outperforming BOLD-fMRI) due to differing inter-modality specificity and resolution. Differing group ID rates, in WF-Ca^2+^ imaging data, as well as clear structure in overall connectome (dis)similarity across groups (and networks), highlights the power of optical imaging measures – specifically at the ‘mesoscale’ – to uncover cell-type specific network structure. We also investigated inter-modal connectome-based ID and found that it did not result in above chance-level ID rates. From an optimistic viewpoint, these observations illustrate the complementarity of WF-Ca^2+^ and BOLD-fMRI and, to an extent, a limitation of coarse static measures like connectome similarity. In-line with these findings, a quantification of within and between mode consistency and differential power^29^, revealed that the most consistent edges are conserved across modes while those with the highest differential power are not.

Altogether, our work affirms using connectome based-ID as a framework for studying individual differences in mouse models. It highlights the power of multimodal methods, that can only be applied in animals, for informing on clinically accessible imaging modes within a framework adopted from the human fMRI literature. Through studying individual differences, across species, neuroimaging can move closer to becoming a means through which to conduct personalized medicine.

## 2. RESULTS

Simultaneous WF-Ca^2+^ and BOLD-fMRI data were acquired from free-breathing lightly anesthetized (0.5-0.75% isoflurane) mice (N = 45, **Table S1**) as we have described previously^30,34^ (**Methods**). To acquire WF-Ca^2+^ imaging data, mice underwent a minimally invasive surgical procedure (7-days prior to the first imaging session, **Methods**) where the skull was exposed and thinned to translucency (but left intact) and a custom glass and plastic head-plate was affixed to the bone for permanent optical access to the cortical surface and immobilization during imaging. All (N = 45) mice underwent three multimodal imaging sessions, with a minimum of 7-days between each session (**Fig. 2A**, left), where structural (for image registration) and functional (BOLD) MRI data were acquired (**Methods**). WF-Ca^2+^ imaging data were acquired simultaneously with the BOLD-fMRI data while mice were in the ‘resting-state’ (i.e., no exogenous stimuli were presented). Forty-minutes of functional multimodal data were acquired at each imaging session across four 10-minute runs. Data were processed using BioImageSuite (BIS) and RABIES (Rodent Automated Bold Improvement of EPI Sequences)^35^ software and registered to an in-house template that has been registered to the Allen Institute Common Coordinate Framework Reference Atlas (CCFv3)^36^ (**Fig. 2B**, **Methods**), as we have described previously^30,34,37^.

**Fig. 2.**
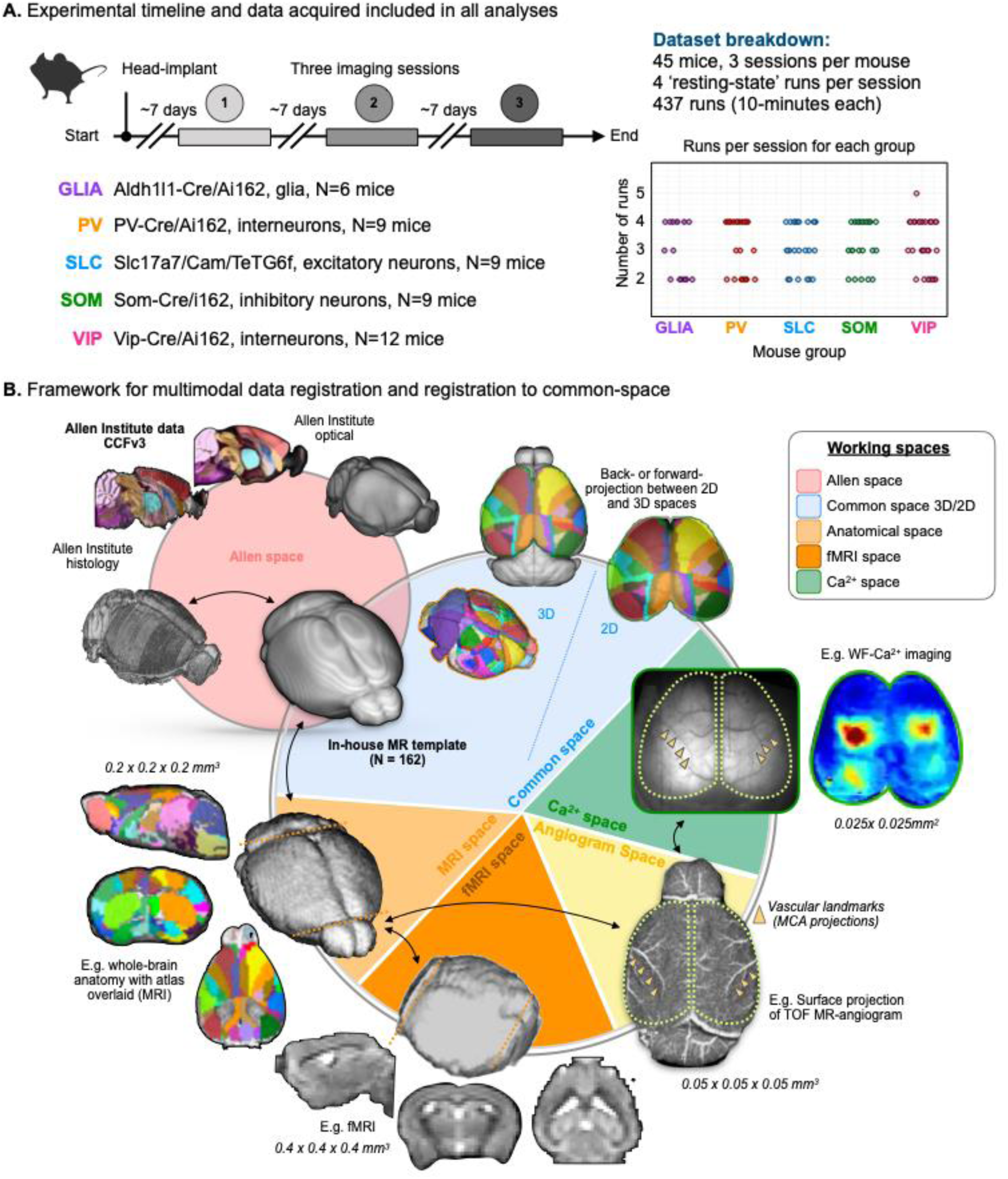
Dataset overview and multimodal data registration framework. **A.** Experiment timeline (left top), group breakdown (left bottom), and number of 10-minute runs included across groups in our analyses (right). **B.** Multimodal registration framework. Structural and functional MR-images (peach and orange), from each mouse at each acquisition (imaging session), were registered to an in-house reference space (**Methods**). This reference space, created from N = 162 datasets collected on our scanner, see ^37^, has been registered to the Allen Institute CCvf3 and Atlas ^36^ (pink). We use an MRI time-of-flight angiogram (yellow) to capture blood vessels on the cortical surface that also appear in the static WF-Ca^2+^ imaging data (green). These serve as anatomical landmarks to link across imaging modalities, as described previously^30^ (**Methods**). Together, these transformations (black arrows) enable us to move atlases, regions of interest, or data from individual to common space and between 2 and 3 dimensions (using forward or backward projection, see ^30^). All transformations were concatenated before being applied. The tools to create all transformations, do projections, the atlases, and our in-house reference space are all freely available (https://bioimagesuiteweb.github.io/webapp/ or ^35^).

Standard data inclusion criteria were applied (**Methods** and full dataset breakdown **Table S1**). We also required that a minimum of 20-minutes of functional data (i.e., two runs) were retained per mouse per session (**Fig. 2A**, right & **Table S1**). Data were matched across modalities such that all the WF-Ca^2+^ and BOLD-fMRI data in our analyses were temporally matched. For WF-Ca^2+^ imaging data, the majority of the power tended to be concentrated in the infra-slow range; which aligns with previous work^38^ (**Fig. S1; Table S2 and S3**). Hereon, we explored a slow and a fast WF-Ca^2+^ imaging frequency band: (1) 0.008-0.2Hz, WF-Ca^2+^ and (2) 0.4-4Hz, WF-Ca^2+^, as we have done previously^38^. The WF-Ca^2+^ band was chosen to match the BOLD-fMRI data.

Connectomes were computed (using frequency filtered data), using the product-moment correlation (Pearson correlation followed by a Fisher-Z transform), for the whole-brain (BOLD-fMRI only) and cortex (BOLD-fMRI and WF-Ca^2+^ imaging data) as the WF-Ca^2+^ imaging has a limited field-of-view (FOV). Thus, we consider data in four ‘conditions’: (1) WF-Ca^2+^, (2) WF-Ca^2+^, (3) BOLD-fMRI cortex-only, and (4) BOLD-fMRI whole-brain (see **Terminology, Supplementary Methods**). For the BOLD-fMRI whole-brain condition, we used 172 regions from the Allen Atlas (**Fig. S2**). Forty-four of these were ‘visible’ from above (in all N = 45 mice at all sessions) and comprise the regions within our cortical connectomes or remaining three ‘conditions’. The multimodal data registration pipeline and images of both the whole-brain and cortex-only atlases are shown in **Fig. 2B**. In all subsequent figures we indicate the imaging modality, FOV (whole-brain or cortex), and ‘condition’ using black-and-white icons.

### 2.1 Average connectomes showed group and condition-specific features

We investigated differences between functional connectomes within each group to establish whether any genotype-specific characteristics underpin connectome structure. Connectomes were computed at the level of each run (i.e., using 10-minutes of WF-Ca^2+^ or BOLD-fMRI data) with data residing within our in-house reference space (**Fig. 2**). After data-inclusion criteria were applied (**Methods**), we retained N = 437 connectomes per condition. All following analyses were performed on these N = 437 connectomes as well as the within-session, within mouse, averaged connectomes (i.e., averaging across 2-4 run-level connectomes per mouse per session which yielded N = 135 connectomes). None of our findings were affected by using run-level or averaged connectomes. As an overview of these data, we computed ‘grand average’ connectomes (mean across groups, sessions, mice, and runs) for each condition (WF-Ca^2+^, Slow/Fast and BOLD, cortex/whole-brain) (**Fig. 3A**). We also computed average connectomes for each group (**Fig. S3A)**. The standard deviations (SD) of the group-specific average connectomes from the grand average (**Fig. 3A**), for each condition, are shown in **Fig. S3B**.

**Fig. 3.**
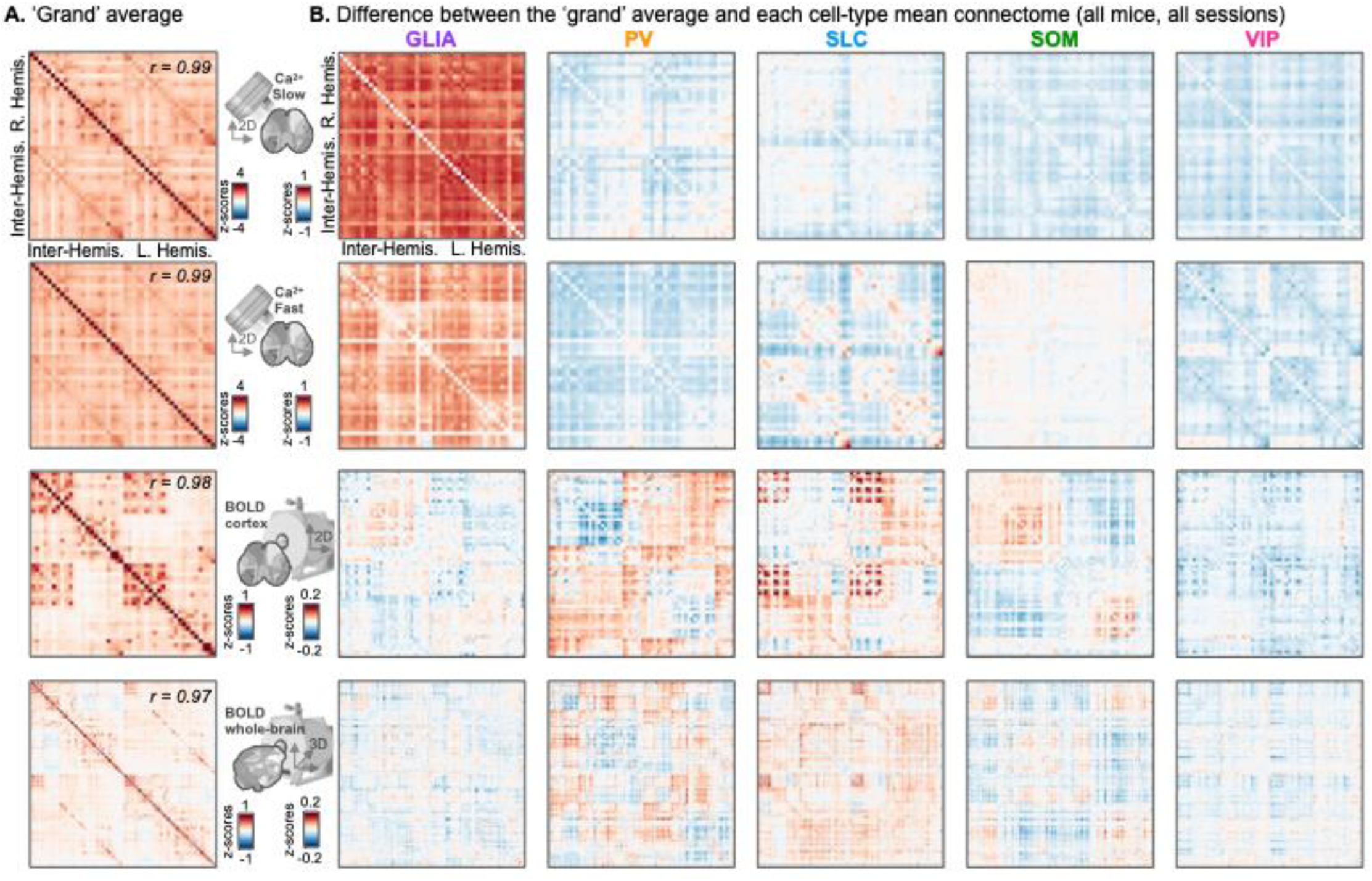
‘Grand’ average connectomes within each condition, and the difference between each group and the grand average (within condition). **A.** ‘Grand’ average connectomes within each condition (averaged across groups, mice, runs, and sessions). Black-and-white icons indicate each condition (top-to-bottom: WF-Ca^2+^_Slow_, WF-Ca^2+^_Fast_, BOLD-fMRI cortex, and BOLD-fMRI whole-brain. Spearman’s correlation, to assess bilateral symmetry reported with r values and p<0.00001 for all conditions. **B.** Group differences between ‘grand’ averages (in **A.**) and each group average (within condition). Number of contributing runs to the dataset average was held constant across groups (N = 56 per group). Groups, left-to-right: GLIA, PV, SLC, SOM, and VIP.

All conditions showed clear bilateral symmetry which indicated greater connectivity between homotopic regions (**Fig. 3A**). To highlight inter-group differences, within condition, the difference between the grand average (**Fig. 3A**) and each group (cell-type) average (**Fig. S3A**) was computed (**Fig. 3B**). For this calculation, the number of runs included from each group was held constant (N = 56 per group).

Group specific differences were found across conditions (**Supplemental File Stats**). For example, in the WF-Ca^2+^ condition, groups (cell-type) exhibited differences in FC strength (*F*(4,4725) = 672.8, p < 0.00001). Post-hoc multiple comparisons (Tukey-Kramer test; **Supplemental File Stats**) indicated that GLIA exhibited the highest FC strength relative to all other groups (e.g., GLIA vs. PV mean difference = 0.61, 95% CI [0.57, 0.66], p < 0.00001). Among interneuron subtypes, VIP showed significantly lower FC than PV (mean difference = 0.08, 95% CI [0.03, 0.12], p < 0.00001) and no difference from SOM (p = 0.34). Further PV did not differ from SLC (p = 0.5). SLC exhibited higher FC than SOM and VIP (e.g., SOM vs. SLC mean difference = −0.07, 95% CI [−0.12, −0.03], p < 0.0001). These findings highlight distinct FC patterns among groups, with GLIA showing the highest FC strengths.

For WF-Ca^2+^, we observed differences in FC across groups (*F*(4,4725) = 399.2, p <0.00001), with post-hoc testing showing a similar pattern as observed for WF-Ca^2+^ (**Supplemental File Stats**). The BOLD-cortex data showed more inter-group features than expected, although their effects were smaller when compared to the WF-Ca^2+^ data (**Supplemental File Stats;** *F*(4,4725) = 11.58, p <0.00001): opposing patterns between PV and SOM (**Fig. 3B**) (p < 0.00001), no differences between GLIA and SOM, while SLC had higher FC than both SOM and VIP (p<0.00001).

### 2.2 Inter-connectome, intra- and inter-condition correlations showed group and condition-specific features

Inter-connectome correlations were computed within (e.g., WF-Ca^2+^_Fast_ to WF-Ca^2+^_Fast_) and between (e.g., WF-Ca^2+^_Fast_ to BOLD-fMRI) conditions. Within condition comparisons were computed across time (e.g. WF-Ca^2+^ session 1 to WF-Ca^2+^ session 2), while between condition comparisons were computed within session using simultaneously acquired multimodal data (e.g. WF-Ca^2+^ session 1 to BOLD cortex session 1) (**Fig. 4** & **S4**). WF-Ca^2+^ connectomes were compared within matched frequency bands (Slow and Fast in **Fig. 4A** & **B**, and **Fig. S4A** & **B**, respectively) and to BOLD-fMRI cortex-only data. Comparisons were computed for all six possible session pairings: session 1 « 2 (**Fig. 4**), 1 « 3 (**Fig. S4, left**) and 2 « 3 (**Fig. S4, right**). Data were organized by group (top-to-bottom: GLIA, PV, SLC, SOM, & VIP) to highlight between group features. For session 1 « 2, within condition (WF-Ca^2+^), an effect of group on FC similarity (*F*(4,4341) = 524.6, p < 0.00001), as well as session interaction (*F*(4,21015) = 657.6, p < 0.00001) was observed. For a breakdown of all between-groups (and sessions) comparisons see **Supplemental File Stats**. Specifically, GLIA (topmost) and VIP (bottommost) showed lower within and between condition inter-connectome correlation strengths relative to other groups for the WF-Ca^2+^ condition (**Fig. 4B**), as shown by multiple comparisons analyses (e.g., GLIA vs. PV mean difference = −0.18, 95% CI [−0.19, −0.16], p < 0.00001; SLC vs. VIP mean difference = 0.19, 95% CI [0.18, 0.20], p < 0.00001). Further, correlation magnitude was on average stronger within WF-Ca^2+^ signals than BOLD-fMRI (**Supplemental File Stats**).

**Fig. 4.**
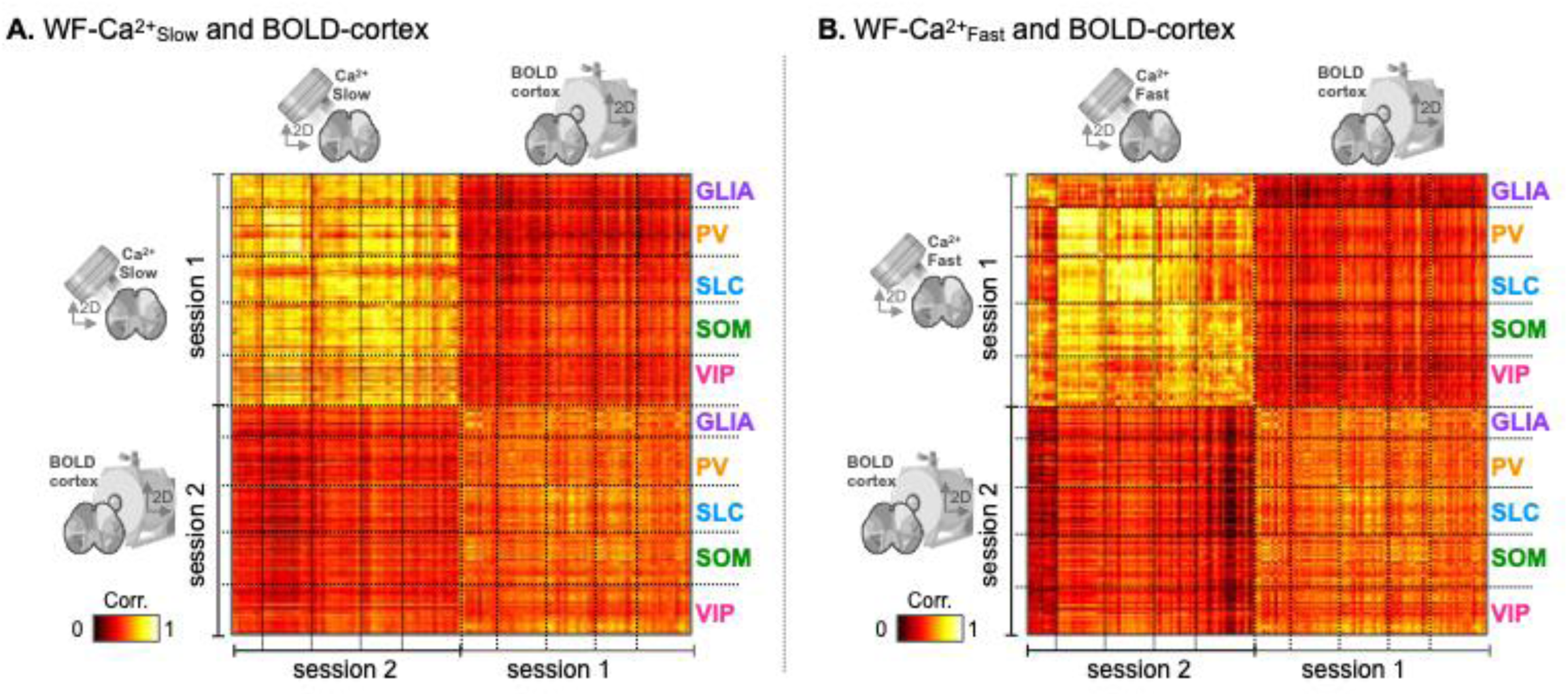
Within and between condition inter-connectome correlation. **A.** WF-Ca^2+^_Slow_ data and BOLD-fMRI cortex-only data. **B.** WF-Ca^2+^_Fast_ data and BOLD-fMRI cortex-only data. Inter-connectome correlation strengths within (upper left and lower right quadrants) and between (upper right and lower left quadrants) conditions using data from within or between sessions, respectively, such that within condition comparisons were across time (i.e., from different sessions), and between condition comparisons used simultaneously collected multimodal data. Each row/column in the matrices is a single run. Session 1 « 2 pairings are shown. For session 2 « 3 & 1 « 3 pairings, see **Fig. S4**. Inter-connectome comparisons were organized by group (top-to-bottom: GLIA, PV, SLC, SOM, & VIP) as indicated.

### 2.3 All intra-condition ID rates of mouse and group were above chance-level with WF-Ca^2+^ outperforming BOLD-fMRI and WF-Ca^2+^ outperforming WF-Ca^2+^

For intra-condition ID analyses, of individual mouse (**Fig. 5A**) or group (**Fig. 5B**), all six possible inter-session pairings were considered. A correct ID was a binary decision based on whether the connectomes that were most similar (highest inter-connectome correlation strength) between sessions belonged to the same mouse or group. For group ID, the mouse being queried was removed from the target session to avoid bleed-through of mouse ID into group ID. As in our analysis of human BOLD-fMRI data (**Fig. 1**), subsets of randomly selected nodes were used to perform mouse or group ID (N = 50 iterations, colored dots in **Fig. 5A** & **B**). These ranged from 5 to all nodes in our cortex-only (N = 44) or whole-brain (N = 172) atlases. Statistically significant ID rates were computed relative to a null distribution generated by shuffling mouse or group ID labels in the target session (N = 50 iterations, greyscale dots in **Fig. 5A** & **B**).

**Fig. 5.**
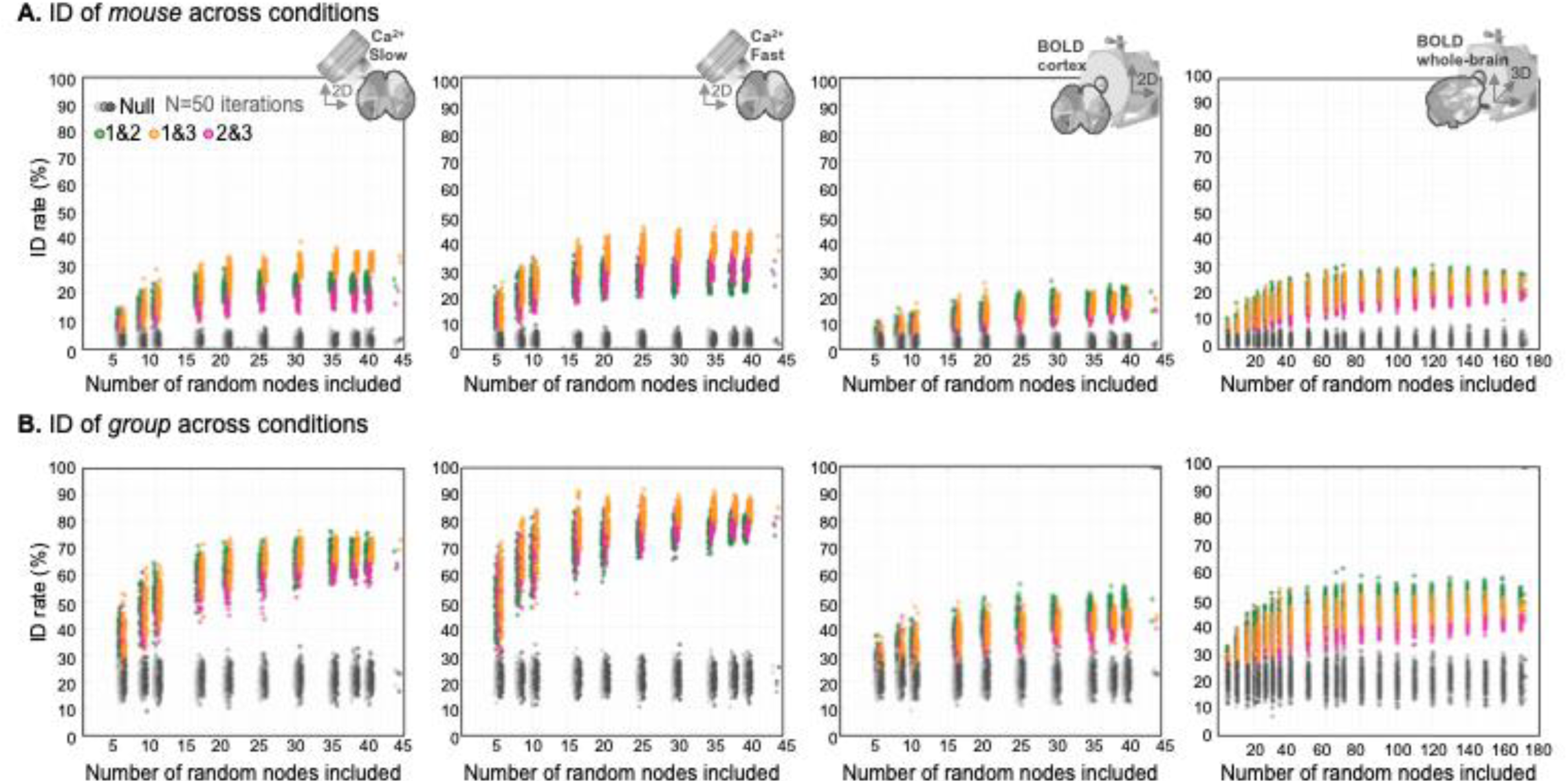
Intra-condition mouse and group ID. **A.** Mouse ID rates for all conditions (left-to-right: WF-Ca^2+^_Slow_ WF-Ca^2+^_Fast_, BOLD-fMRI cortex-only, and BOLD-fMRI whole-brain). **B.** Group ID rates for all conditions (ordered as in **A.**). In **A.** and **B.**, colored dots (green, yellow, & pink) – from intact data – were compared to a null distribution (greyscale dots) where the mouse or group ID labels in the target session were shuffled (N = 50 iterations). For group ID (**B.**), the mouse being queried was removed from the target session (to avoid mouse ID bleed-through). As in Fig. 1, mouse or group ID was repeated using randomly selected nodes ranging from 5 to all-nodes within the atlas (N = 50 iterations). In all cases, intact data outperformed shuffled data.

To investigate ID rates within each condition, all session pairings were compared to a null distribution (MATLAB, *ttest2* Bonferroni corrected). All intra-condition mouse and group ID rates quickly rose above chance-levels (**Fig. 5A** & **B**). Across all conditions, only five nodes were needed for significant ID rates (p < 0.00001 for all conditions, see **Supplemental File Stats**). Similarly, in all cases, correct mouse and group ID rates appeared to reach a plateau when approximately half of all available nodes were being considered (based on a visual inspection of the results, **Fig. 5A** & **B**). When we investigated the apparent session pairing dependence of mouse and group ID rates (in WF-Ca^2+^ data), where longer inter-session times appear to perform better than shorter inter-session times, we found these effects to be significant (MATLAB, *anovan*, factor 1 being the reference session, factor 2 being the target session, multcompare, see **Supplemental File Stats**). There was also an effect of condition on mouse (*F*(2,897) = 622, p < 0.00001) and group (*F*(2,897) = 6046.5, p < 0.00001) ID rates, with WF-Ca^2+^_Fast_ outperforming both WF-Ca^2+^_Slow_ and BOLD-cortex (e.g., in ID of mouse, WF-Ca^2+^_Fast_ vs. BOLD-cortex, mean difference = 14.6, 95% CI [13.6, 15.57], p < 0.00001) and WF-Ca^2+^_Slow_ outperforming BOLD-cortex (mean difference = 8.13, 95% CI [7.16, 9.1], p< 0.00001). Although group ID rates based on WF-Ca^2+^ imaging data were substantially greater than those based on BOLD-fMRI data (**Fig. 5**), the still clearly significant group ID rates achieved with BOLD-fMRI data were noteworthy (**Fig. 5B**). Finally, differences in mouse and group ID rates between cortex-only and whole-brain BOLD-fMRI data were small (despite still being significant for 60 or more nodes, p<3.5e^-5^, MATLAB *ttest2*) which indicated that subcortical areas played a limited role in ID success rate (as anticipated based on work in human subjects, **Fig. 1**).

For both mouse and group, we computed whether correct IDs were shared across conditions (**Fig. S5**). For example, if mouse-1 was correctly matched in the WF-Ca^2+^ calculation (leftmost panel, **Fig. 5A**), was mouse-1 also correctly matched in the BOLD-fMRI whole-brain calculation (rightmost panel, **Fig. 5A**)? Both intra- and inter-modality comparisons were compared to a null distribution generated by shuffling the correct hit labels from one of the two calculations (N = 50 iterations). Overall, correct IDs of mouse and group were not shared when results from different modalities were compared (**Fig. S5 A** & **B**). Conversely, correct IDs of mouse and group showed consistent above chance-level subject overlap within modality (**Fig. S5 C** & **D**).

### 2.4 Intra-condition ID rates of mouse and group showed modest *a priori* functional network dependence

Human BOLD-fMRI data-based connectome ID shows a strong dependence on *a priori* functional networks (**Fig. 1**)^5,7^. To search for parallels in mouse data, we investigated 12 whole-brain (BOLD-fMRI) *a priori* canonical networks (somatosensory, auditory, visual, olfactory, sensory, cortex, limbic, lateral cortical, DMN, salience, subcortical, and brainstem). Six of these (somatosensory, visual, sensory, limbic, lateral cortical, and DMN) were ‘visible’ from above (**Fig. S2**) and used in our WF-Ca^2+^ imaging and BOLD-fMRI cortex-only analyses.

As in **Fig. 1**, we determined whether canonical networks played a disproportionate role in mouse or group ID by comparing ID rates obtained when a network was used to random size-matched selections of nodes (N = 50 iterations) (**Fig. S6 A** & **B**). For WF-Ca^2+^ imaging data, none of the canonical networks performed better than this benchmark for mouse or group ID. Somewhat surprisingly, several canonical networks under-performed for group ID (**Fig. S6B**). Conversely, the BOLD-fMRI data showed some parallels to results obtained using human BOLD-fMRI data. The whole-brain lateral cortical network (a proxy for the human frontal parietal network) and somatosensory network produced greater mouse ID rates while the subcortical and brainstem networks under-performed (**Fig. S6A**). Notably, the nodes in the lateral cortical and somatosensory networks that were ‘visible’ from above and were thus included in our BOLD-fMRI cortex-only analysis, did not produce the same results as in our whole-brain analyses. These observations suggest that connections to regions outside of the WF-Ca^2+^ imaging FOV may play a disproportionately important role in increasing mouse ID rates and could help explain why we did not see similar results in the WF-Ca^2+^ imaging data.

Overall, given the notable role played by canonical networks in human neuroimaging studies (see **Fig. 1**), these results indicate that data collected from mice do not recapitulate this strong relationship between established networks and ID rates. This could be due to inter-special differences (i.e., mice lack the higher-order brain circuits that perform well in human-based ID), or that canonical networks in mice are less well-defined.

### 2.5 Intra-condition ID rates depended on *a priori* canonical network and group (e.g., from WF-Ca^2+^)

The preceding subsection considered all groups together. Here, we searched for differences in mouse ID rates across groups (e.g., are SLC more identifiable than GLIA?) and structure within the incorrect IDs. For example, are GLIA that were incorrectly identified most similar to others in the GLIA group or are they frequently misidentified as members of another group (e.g., PV)? For this aim, we interrogated a range of thresholds for “top-hits” (i.e., most similar inter-session intra-contrast connectomes) in-place of selecting only one top-hit (as done in the preceding subsections). We identified the top 5, 15, or 25 most correlated inter-session connectome pairs (when considering runs, N = 146/144/147 for sessions 1/2/3), and the top 2, 4, or 6 most correlated inter-session connectome pairs (when considering averaged runs, N = 45 per session). An example is shown in **Fig. 6A** for WF-Ca^2+^ (sessions 1 & 2), all nodes. We quantified the binarized “top-hit” matrices (**Fig. 6A**, right) as the fraction of hits within each area of interest where the areas of interest were each within- or between-group pairing (e.g., GLIA-to-GLIA, indicated in **Fig. 6**). The fraction of top-hits within each area of interest was expressed as a matrix (**Fig. 6B**) for all session pairs, thresholds, conditions, and canonical networks. As above, a null distribution was generated by shuffling the target session labels (N = 50 iterations).

**Fig. 6.**
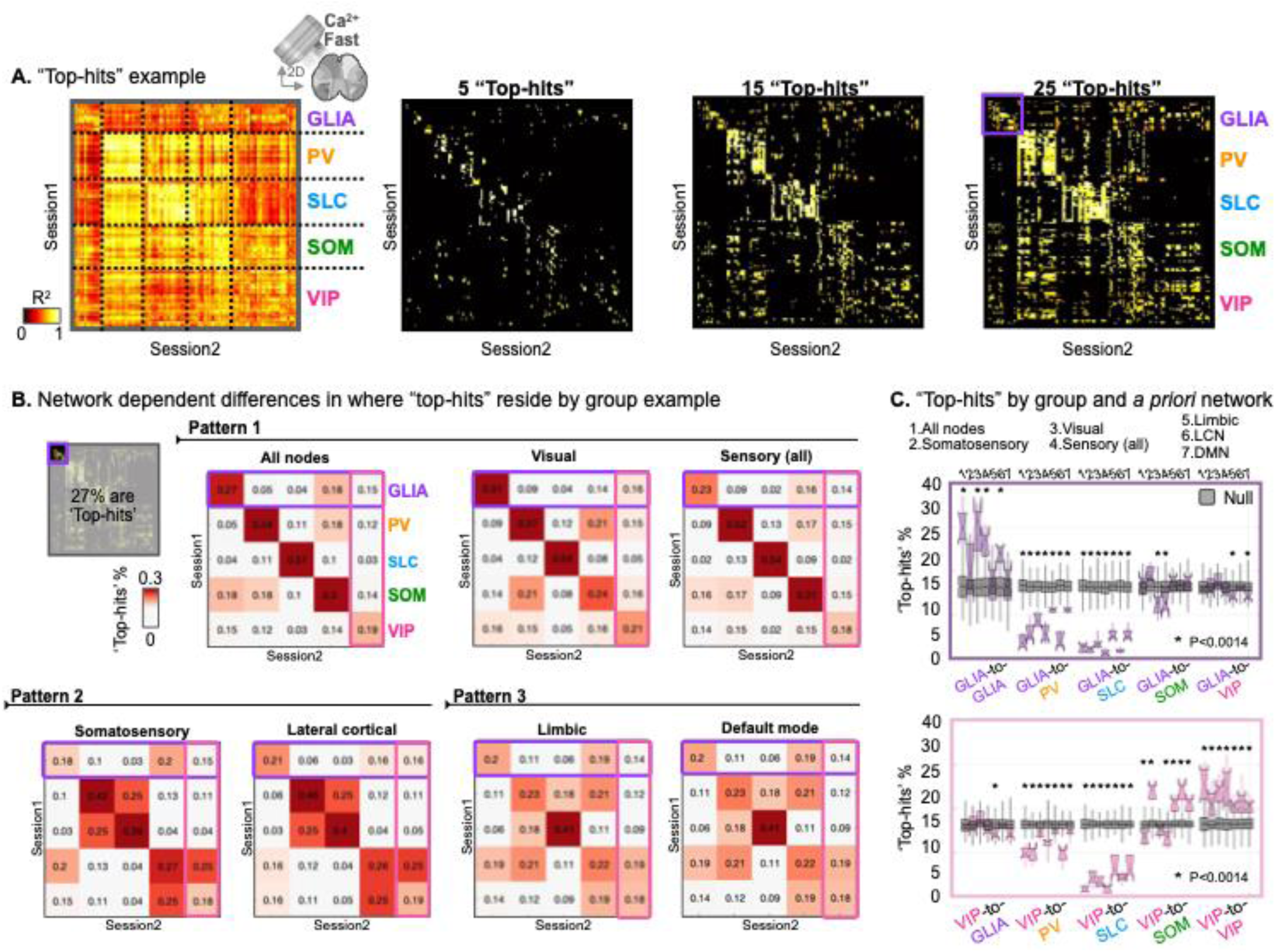
Representative example of network influence on intra-condition (WF-Ca^2+^ _ast_) ID separated by group. **A.** Left-to-right. Intra-condition (WF-Ca^2+^_Fast_) connectome-to-connectome correlation matrix for sessions 1 & 2. Leftmost matrix (sans threshold) was replicated from Fig. 4**. B**. Rightmost three matrices are with thresholds (5, 15, & 25) applied. The purple square indicates the area of interest where GLIA-to-GLIA matches were located (corresponds to the purple square in **B.**). **B.** Fraction of ‘top-hits’ within each area of interest (within and between group pairings) for WF-Ca^2+^_Fast_, sessions 1 & 2, all runs, threshold 25, summarized “top-hit” fraction matrices. Results for all nodes (N = 44), as well as six canonical networks (‘visible’ from above), see **Fig. S2**. Matrices are arranged into three patterns. Color-scale was adjusted such that the red hue indicates a value greater than the null distribution (generated by shuffling target session labels, N = 50 iterations) **C.** Example boxplots for GLIA (top, purple), and VIP (bottom, pink) results binned left-to-right by group pairing. Within each bin, results from each a priori network are plotted. Distributions were generated from the six session pairings. Thus, one data point per distribution, in **C.**, corresponds to one row/column in **B.**, circled in purple (GLIA) or pink (VIP). As before, null distributions (grey) were generated by shuffling target session labels (N = 50 iterations). Group pairings where a statistically significant (MATLAB ttest2, Bonferroni corrected) surplus, or paucity, of ‘top-hits’ were found are indicated by an *.

The example matrices in **Fig. 6B** were organized into three clusters to highlight three patterns. First, all nodes, visual and sensory networks (top row) showed a strong diagonal which indicated high within-group ID rates (although, VIP performed poorly relative to the rest) and low between group incorrect IDs. The second pattern (bottom row, left), evident in the somatosensory and lateral cortical networks, showed low GLIA ID rates (GLIA being incorrectly identified as SOM), reasonable PV, SLC, and SOM ID rates – albeit with frequent errors between PV and SLC as well as between SOM and VIP – and, as in the first pattern, low correct VIP ID rates. In the final pattern, limbic and DMN, showed low (but still significant) within-group ID rates with more diffuse errors (than seen in the second pattern). Results were summarized further, across session pairs and canonical networks, using boxplots (**Fig. 6C**).

The example results in **Fig. 6C** (all runs, threshold 25, WF-Ca^2+^, GLIA and VIP) showed that where ‘top-hits’ were found, had group-pairing and network dependence. For instance, limbic and default mode networks, showed poor correct GLIA ID rates (same as chance) (**Fig. 6C**, top). Conversely, VIP (**Fig. 6C**, bottom) showed more modest correct ID rates without clear differences between networks. In terms of where incorrect IDs resided, GLIA were unlikely to be misidentified as PV or SLC (below chance) but were mistaken for SOM or VIP at chance-level (mostly regardless of network). Similarly, VIP were unlikely to be misidentified as SLC, or PV – albeit to a lesser extent – and were mistaken for GLIA at chance-level. VIP were, however, misidentified as SOM at greater than chance-level (with some network dependence). These observations, and others for remaining groups, held across ‘top-hit’ thresholds, and the averaging of runs.

### 2.6 Groups showed some consistent cross-condition network and inter-group ID patterns

The analyses outlined in **subsection 2.5** were performed on all groups using data from all conditions. Representative results (from all four conditions) for GLIA, SLC, and VIP are shown in **Fig. 7**. Results from PV and SOM are shown in (**Fig. S7**). WF-Ca^2+^ and WF-Ca^2+^ conditions showed nearly identical results across groups and canonical networks. SLC, followed closely by PV, showed the highest fraction of correct IDs with GLIA, SOM, and finally VIP showing progressively lower (but still above chance) fractions of correct IDs. Overall, limbic and default mode networks performed poorly, while all nodes and the sensory network performed best. SOM and PV were more likely to be misidentified for one another than either were to be misidentified as SLC. Similarly, PV and SLC showed some propensity for being misidentified for one another; specifically, when the somatosensory and lateral cortical networks were being considered. Finally, it was very unlikely for GLIA to be misidentified as PV or SLC (and vice versa).

**Fig 7.**
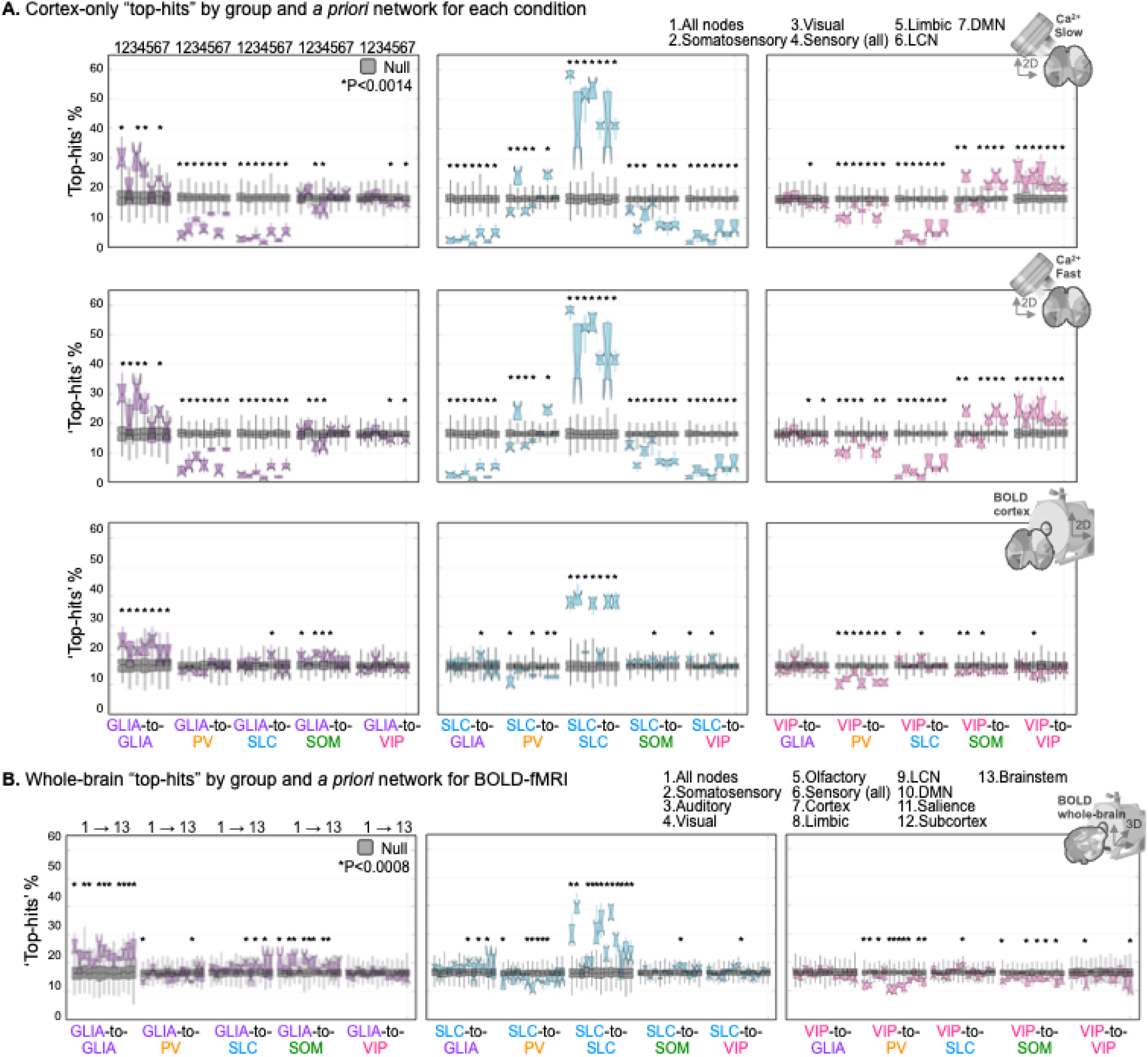
Network influence on intra-condition ID separated by group (for GLIA, SLC, and VIP). Boxplots are constructed as described in **Fig. 6 C.** (topmost left and right panels were replicated from **Fig. 6 C.**). Results for PV and SOM are shown in **Fig. S7**. **A.** Results (top-to-bottom) for WF-Ca^2+^_Slow_, WF-Ca^2+^_Fast_, and BOLD-fMRI cortex-only. Six canonical networks, in addition to all ROIs. **B.** Results for BOLD-fMRI whole-brain. Twelve canonical networks, in addition to all ROIs. For **A.** and **B.** group pairings where a statistically significant (MATLAB ttest2, Bonferroni corrected) surplus, or paucity, of ‘top-hits’ were found were indicated by an *.

Cortex-only BOLD-fMRI results mirrored some features that were evident in the WF-Ca^2+^ imaging data. As in the WF-Ca^2+^ imaging conditions, SLC showed the highest fraction of correct IDs. Although PV were still above chance, they performed more poorly, and not as well as GLIA. SOM were still above chance-level, but VIP were not. These findings roughly paralleled what was seen in the WF-Ca^2+^ imaging conditions. Network effects in the cortex-only BOLD-fMRI condition, were only evident in SLC, and showed that visual and limbic networks performed poorly. Similar patterns, with more clear network dependence, were evident in the whole-brain BOLD-fMRI condition where we considered 12 canonical networks (and all nodes). Here, auditory, visual, and olfactory performed poorly, as did limbic, default mode, salience, subcortical, and brainstem, relative to somatosensory, sensory, cortex, the lateral cortical network, and all nodes. Findings were consistent across thresholds and the averaging of runs.

### 2.7 Cross-modality instances of high connectome-to-connectome correlation strengths showed group and network dependence

Overall, our findings were consistent with our expectations given the appearance of the matrices in **Fig. 4** and **Fig. S4**. Inter-condition comparisons (top left, and bottom right quadrants) showed the classic diagonal structure which indicated that the highest connectome-to-connectome correlations were (on average) between the same animals. The block diagonal structure also indicated that within-group errors were likely to be more frequent than between-group errors. This was abundantly clear when a threshold was applied (**Fig. 6A**). Notably, the same pattern was not evident when connectomes were compared across imaging modalities (top right, and bottom left quadrants in **Fig. 4** and **Fig. S4**), which led to the unsurprising result that when we performed cross-modality ID (BOLD-fMRI cortex-only to WF-Ca^2+^ or WF-Ca^2+^), using the whole dataset in a ‘winner-take-all’ framework we found no significant results (data not shown). Nevertheless, we pursued a more subtle interrogation of cross-modality similarity quantification using a framework that was similar to the one applied in subsections **2.5** & **2.6**.

We took the average of the “forward” and “backward” (e.g., session 1→2, and 2→1) connectome-to-connectome correlation matrices for each pair of sessions (i.e., top right and bottom left quadrants in **Fig. 4, Fig. S4**). Rather than threshold for “top-hits”, we applied a range of correlation strength thresholds (0.45, 0.40, and 0.55) to assess where the greatest agreement was between imaging modalities in terms of connectivity patterns. Null distributions were generated as before by shuffling the target session labels (N = 50 iterations). As in the previous subsections, we computed the fraction of “hits” within each area of interest for all group pairings and canonical networks.

Results, boxplots in **Fig. S8** for BOLD-fMRI cortex-only → WF-Ca^2+^ or WF-Ca^2+^, were organized differently than boxplots in **Fig. 6**, **7**, and **S7** due to clear differences in correlation values between modalities across canonical networks (see y-axis ranges in **Fig. S8**). For example, limbic and default mode networks showed a much higher fraction of high inter-modal correlations than other canonical networks. Thus, boxplots were organized with canonical networks in rows, and group pairings in columns.

For BOLD-fMRI cortex-only → WF-Ca^2+^ (**Fig. S8A**), GLIA, SOM, and VIP showed no instances where matched groups (e.g., GLIA-to-GLIA) showed higher fractions of high inter-modal correlation values than the null distribution. Conversely, PV and SLC showed select instances where fractions of high inter-modal correlation values were above chance. Curiously, these were apparent within different canonical networks (**Fig. S8**). For BOLD-fMRI cortex-only → WF-Ca^2+^ (**Fig. S8B**), GLIA, PV, and SLC showed no instances where matched groups showed a higher fraction of high inter-modal correlation values than chance whereas SOM and VIP showed some select instances that were. However, the specificity of these occurrences was low (i.e., they appeared coincident with high fractions of high inter-modal correlation values in unmatched group pairings, e.g., **Fig. S8B**, somatosensory, SOM-to-PV, SOM-to-SOM, as well as PV-to-SLC, PV-to-PV, and PV-to-SOM, were all above chance).

### 2.8 Edges with high consistency and differential power – reproducibility, brain module representation, and inter-condition overlap

We investigated edges with high consistency and differential power^5,29,33^. Within condition, edges with high consistency or differential power were well-preserved across session pairings (**Fig. S9**). The reproducible edges with high consistency or differential power for each condition are organized by brain module (as defined by Bergmann et al.)^29^ (**Fig. S9C** & **S10**). As expected, based on human neuroimaging as well as the work by Bergmann et al., edges with high and reproducible consistency resided within brain module (**Fig. S10B**, top). Conversely, edges with high and reproducible differential power resided both within and between brain modules (**Fig. S10B**, bottom). We also examined whether edges with high consistency or differential power were shared across conditions (**Fig. S11**). Edges with high consistency were shared across all three conditions, whilst edges with high differential power were not. This was roughly in-line with our finding that inter-condition ID was not successful (see above).

## 3. DISCUSSION

We investigated functional connectome-based ID in mice using simultaneously acquired WF-Ca^2+^ and BOLD- fMRI data. In-line with previous work in mice^29^ and humans^5–16^, animals were identifiable using BOLD-fMRI data. Building on this foundation and affirming the likely cellular-basis of these findings, animals were also identifiable using WF-Ca^2+^ imaging data. To the best of our knowledge, this is the first study to investigate connectome-based ID using this modality. Further, the unique structure of our dataset allowed us to explore group ID rates (where groups bore GCaMP fluorescent labels in different cell populations), as well as inter-modal connectome-based ID for the first time.

Translational work across species can be aided by using common tools (e.g., imaging modalities) and analytical frameworks^39,40^ (here, connectome-based ID). Work of this nature contributes by establishing whether (and the degree to which) specific measurements are possible in multiple species^41^ and by extension how useful animal models might be within a specific analytical framework or clinical context. Given a foothold, i.e., when overlap across species can be established, work in animals can offer insights that are not attainable from human subjects^21^.

Connectome-based ID, using human BOLD-fMRI data, produces high rates of correct IDs (typically about 60-99%, depending on sample size^5,8,13^; here, 88% in the human data, **Fig. 1**). Previous work on mouse BOLD-fMRI data, by Bergmann et al., established that ID rates in mice are lower (60-70%)^29^ but still well above chance. We observed even lower, but again still significant, ID rates (20-40%) using BOLD-fMRI data. As expected, ID rates based on WF-Ca^2+^ imaging data are higher (30-50%), but still below what Bergmann et al. observed. This gap was not addressed by repeating our analyses using a framework designed to match the one implemented in their paper. Specifically, creating a query and target set by splitting the dataset within each session, i.e. using runs from different timepoints such that each timepoint was represented in each averaged connectome (or query and target set) (**Fig. S12 A** & **B**). The remaining notable differences between the study by Bergmann et al. and our own is that our mice were lightly anesthetized (0.5-0.75% isoflurane) whilst theirs were awake, and dataset size (their N = 16 vs. our N = 45; which has previously been shown to affect ID rates in human participants^13^).

To investigate the role played by sample size in mice, we randomly selected N=15 mice (50 iterations) and repeated our mouse and group ID analyses (**Fig. S12 C** & **D**) using the same split-session framework as above. N=15 (in place of N=16) was used so that groups were equally represented (N=3 per group across all iterations). For all runs, mouse ID rates were 36 ± 11% (mean ± SD) with a maximum of 60% when runs were considered. For averaged runs, mouse ID rates were 42 ± 13% with a maximum of 73%. Likewise, for all runs, group ID rates were 50 ± 9% with a maximum of 70%, when runs were considered. For averaged runs, group ID rates were 28 ± 10% with a maximum of 67%. Overall, these results affirm that smaller sample sizes produce higher ID rates (for mouse and group ID), and that sample size likely contributed to the difference in ID rate reported by Bergmann et al. and within the present work.

Similarly, it is likely that our use of anesthesia reduced our observed ID rates given recent work in anesthetized human subjects^42^. This is notable given that the majority of fMRI studies in rodents use anesthesia, and that differences in resting-state fMRI signals between awake and anesthetized mice have not been extensively characterized (see ^43^ for a recent systematic review). Our use of anesthesia also likely contributed to some of the differences in edges with high consistency and differential power that we and Bergmann et al. report. The inherent challenges (e.g., motion and stress), and significant time required, to perform fMRI experiments on awake mice necessitates clear and specific scientific reasons for doing so. It is also exceptionally difficult, and resource intensive, to generate large datasets (typical of ID-based or similar analyses) from awake mice^44^. Future work is needed to confirm whether the use of anesthesia in mice truly reduces connectome-based ID rates and the role this may play in studies focused on inter- and intra-subject differences. In combination with follow-up studies, the findings in this work may provide some key evidence for using awake animals within the context of exploring individual differences.

Human BOLD-fMRI data-based connectome ID shows clear canonical network dependence (**Fig. 1**)^1,23,25^. This feature was not evident in the study by Bergmann et al. where ID rates were comparable between all cortical nodes, association, and sensory systems. Here, we extended this analysis using a broader range of networks, including subcortical areas, and found some modest indication that networks may influence ID rate. Specifically, paralleling findings from human BOLD-fMRI data, we found somatosensory, and lateral cortical networks performed relatively well (for their size), whilst subcortical and brainstem networks performed poorly (**Fig. S6**). Notably, data from the whole-brain was necessary for this observation as these findings were not evident in cortex-only BOLD-fMRI or WF-Ca^2+^ imaging data. Given the overall low ID rates in our dataset and the absence of higher-order association networks—believed to be key drivers of individual differences in humans—it is unsurprising that these effects were modest^45,46^.

Our unique multimodal dataset allowed us to explore new aspects of the connectome-based ID framework. With our WF-Ca^2+^ imaging data, we showed that the framework extends beyond applications in BOLD-fMRI data, which helps affirm that cellular activity, are the likely drivers of correct IDs. Although we did not observe cross-modal agreement in terms of which mice are correctly identified across all comparisons, we did see significant agreement between WF-Ca^2+^ and cortex-only BOLD-fMRI (**Fig. S5A**) – the most closely matched cross-modal conditions. However, our finding that reproducible edges with high differential power (**Fig. S11**) were not shared across conditions is further evidence that there are complexities that warrant further investigation.

Our multimodal data also allowed us to explore group ID, which was well above chance using data from either modality. The high rates from the WF-Ca^2+^ imaging data indicated that there are differences in connectivity structure that are apparent in data from different cellular sources at our ‘mesoscale’ spatiotemporal resolution – even at low frequencies. This is encouraging for future work aimed at examining cell-type specific activity or network-based measures at the ‘mesoscale’ using WF-Ca^2+^ imaging data. This finding runs counter to the assumption that if cells of different types are closely linked at the microscale, their apparent meso- or macro-scale organizational structures will be indistinguishable.

We also used these data to characterize inter-group connectome similarity and dissimilarity (**Fig. 6**, **7**, & **S7**). Clear patterns in the WF-Ca^2+^ imaging results emerged between groups that aligned with how closely related different cell-types are at the micro-circuit level. For example, GLIA were more easily dissociated from PV and SLC, whilst the later were more similar to one another. Similarly, SOM and VIP were similar to one another and unlike SLC. The broad similarities between inhibitory interneurons (and differences with respect to excitatory neurons and glial cells) generally aligns with previous WF-Ca^2+^ work^38^ and is consistent with the hierarchical nature of cell type classes^47^. Curiously, some of these patterns were recapitulated in the fMRI data where the signal source (BOLD) should be independent from group membership except for the role played by closer lineage. Specifically, we expected that mice might be more similar within group, but not that trends across groups would emerge (e.g., for SLC to be the most identifiable, and for VIP to be at chance-level). This could indicate that there are differences across groups (e.g., sensitivity to anesthesia) that contributed to some of the observed inter-group differences in both modalities. Future work in awake mice, and animals bearing more than one fluorescent cell-label are needed to investigate this finding further. Yet, the difference in effect size, as well as the clear dissimilarities evident in the WF-Ca^2+^ imaging data - but not the BOLD-fMRI data – indicate that the cell-type specific findings in the WF-Ca^2+^ imaging data are still likely present.

Finally, we found that a strict winner-take-all ID framework failed when applied across imaging modalities (i.e., WF-Ca^2+^ → BOLD-fMRI and vice versa). On the one hand, this outcome emphasizes the complementary information available from each modality and encourages future experiments that investigate how these contrasts, when applied together, can further inform on brain function^48^. On the other hand, given that WF-Ca^2+^ imaging is a more proximal measure of neural activity than BOLD-fMRI, this could fuel skepticism for the neural-basis of BOLD-fMRI-based measures of functional connectivity. Our finding that edges with high consistency are reproducible and shared across conditions whilst reproducible edges with high differential power are not shared across conditions is in-line with there being some truth to both these arguments.

To investigate further, we examined whether different groups, or networks, showed greater connectome similarity across modes than would be expected by chance (**Fig. S8**). We found some preliminary, but intriguing evidence that SLC and PV groups, showed greater inter-modal similarity (albeit unexpectedly using WF-Ca^2+^_Fast_ whilst some, more tenuous patterns, were found for SOM and VIP using WF-Ca^2+^_Fast_). More in-depth investigations into these data are warranted and necessitates a framework that goes beyond measures of connectome similarity. Future work could also use data obtained via other imaging modalities^49^, as well as examining the extent to which identifiability is affected by disorders commonly modelled in mice (e.g.^50^).

While this study provides new insights into connectome-based identifiability in mice, several limitations should be noted. First, network-level analyses showed mixed results, suggesting that functional networks in the mouse brain may not contribute to ID in the same way as in humans or that they require further development. Future work should explore alternative network definitions and parcellation strategies to clarify their role. Second, our use of light anesthesia likely influenced ID rates. While imaging awake mice (especially at scale) is challenging, future research should determine the extent of this effect. Finally, although our heterogeneous dataset – comprised of mice with five different fluorescent targets – allowed us to investigate some novel aspects of connectome-based ID, it may have obscured some of our analyses.

In conclusion, using a unique simultaneously acquired WF-Ca^2+^ and BOLD-fMRI dataset, we showed that mice are identifiable using a connectome-based ID framework adopted from the human BOLD-fMRI literature. This work extends this framework beyond BOLD-fMRI and explores features of connectome identifiability, similarity, and dissimilarity across subjects, groups, and imaging modalities. Although a strength of animal research is the ability to conduct experiments within a tightly controlled environment and on nearly identical subjects, as neuroimaging methods in mice become more sophisticated, studies of inter- and intra-subject features stand to contribute in new ways. Areas where this may be most relevant include studies of mental illness, or within any arena where heterogeneity – both in terms of brain function and behavior – of unknown origin and significance are the norm^27,51^.

## 4. METHODS

All procedures were performed following the Yale Institutional Animal Care and Use Committee (IACUC) and in agreement with the National Institute of Health Guide for the Care and Use of Laboratory Animals.

### 4.1 Group and dataset overview

Mice were housed on a 12-hour light/dark cycle. Food and water were available *ad libitum*. Animals were 6-8 weeks old, 25-30g, at the time of the first imaging session. Five mouse lines were used: (1) GLIA, (2) PV, (3) SLC, (4) SOM, and (5) VIP (**Fig. 2A** & **Table S1**). All groups were of mixed sex.

Four groups (all but GLIA) shared a C57BL/6J background (**Table S1**). Male CRE mice were selected from the offspring of parents with different genotypes; this avoided leaking of CRE expression. See **Table S1** for the full breeding scheme. Briefly, the Ai162 genotype resulted from tTA and TITL-GCaMP6s (TIGRE1.0). Differences between TIGRE1.0 and 2.0 have been reported^52^ and TIGRE1.0 mice have been previously described^30^. The number of inbred generations may have caused some genetic variation between groups. Specifically, VIP mice were bred in-house for a prolonged period, which may have led to slight variations in their background. Given the careful monitoring of sublines by Jackson’s lab, this factor was likely negligible for other groups. The fifth group (GLIA) was derived from the Aldh1l1-CreER/Ai162 line, where Aldh1l1-CreER is on a C57BL/6;FVB/N genetic background. The GLIA group received tamoxifen injections for three consecutive days after weaning to elicit GCaMP expression. No visible physiological or phenotypic effects due to tamoxifen toxicity have been reported in these animals.

### 4.2 Surgical procedure for permanent optical access to the cortical surface and immobilization

All mice underwent a minimally invasive surgical procedure were an in-house built head-plate (acrylonitrile butadiene styrene plastic, TAZ-5 printer, 0.35mm nozzle, Lulzbot) was attached to the thinned, but intact, skull for permanent optical access to the cortical surface, necessary for WF-Ca^2+^ imaging, and immobilization. The surgical procedure has been described by us previously^30,34,38^. Briefly, mice were initially anesthetized with isoflurane (5%, 70/30 medical air/O_2_) and head-fixed in a stereotaxic frame (KOPF, USA). Isoflurane was then reduced to 2%. An eye ointment was applied; meloxicam (2mg/(kg)body weight) administered subcutaneously and bupivacaine (0.1%) injected locally, at the incision site. After shaving the hair, the scalp was washed with betadine followed by ethanol 70%, (×3). The scalp was surgically removed, and the skull cleaned and dried. Antibiotic powder (Neo-Predef) was applied to the incision site, and isoflurane reduced (1.5%). Skull-thinning of the frontal and parietal skull plates was performed with a hand-held drill (FST), diameter of the tips 1.4mm and 0.7mm. Superglue (Locite) was applied to the exposed skull, followed by transparent dental cement C&B Metabond (Parkell). The head-plate was attached to the dental cement before solidification. The head-plate had a double-dovetail plastic frame with a microscope slide hand-cut to match the size and shape of the mouse skull^19^.

### 4.3 Imaging protocol

Mice were allowed 1-week to recover post-surgery before undergoing three simultaneous WF-Ca^2+^ and BOLD- fMRI sessions on an 11.7T MRI scanner (Bruker, Billerica, MA) using custom MRI-compatible WF-Ca^2+^ imaging equipment^30^. The interval between each imaging sessions was at least 1-week.

#### 4.3.1 BOLD-fMRI data acquisition

Data were acquired as described previously^30,34^ using ParaVision 6.0.1. Mice were scanned whilst lightly anesthetized with isoflurane and free breathing (0.5-0.75%, 70/30 medical air/O_2_). Body temperature was continuously monitored (Spike2, Cambridge Electronic Design Limited), and maintained with a circulating water bath. Imaging data and physiological recordings were synchronized (Master-8 A.M.P.I., Spike2 Cambridge Electronic Design Limited). A gradient-echo, echo-planar-imaging (GE-EPI) sequence with a 1.0s repetition time (TR) and 9.1ms echo time (TE) was used with an isotropic resolution of 0.4×0.4×0.4mm^3^. Twenty-eight slices yielded near whole-brain coverage (**Fig. 2B**). Each functional run was comprised of 600 volumes or 10mins of data. On average, 7 functional runs were acquired per mouse per session. Three included a unilateral LED-stimulation whilst 4 were acquired in the resting-state (no external stimuli were presented). Here, we consider the resting-state data only.

In addition to the functional data, 4 structural images were acquired during each imaging session for data registration purposes^30,34^. (1) A multi-spin-multi-echo (MSME) image sharing the same FOV as the functional data with a TR/TE of 2500/20ms, 28 slices, two averages, and resolution of 0.1×0.1×0.4mm^3^ in 10mins and 40s. (2) A whole-brain isotropic MSME image with a TR/TE of 5500/20ms, 78 slices, two averages, and resolution of 0.2×0.2×0.2mm^3^ in 11mins and 44s. (3) A fast-low-angle-shot (FLASH) time-of-flight (TOF) angiogram, with a TR/TE of 130/4ms, resolution of 0.05×0.05×0.05mm^3^ and FOV of 2.0×1.0×2.5cm^3^ in 18mins. (4) A FLASH image of the angiogram FOV, with a TR/TE of 61/7ms, four averages, and resolution of 0.1×0.1×0.1mm^3^ in 11mins and 24s. Each average of images (1) and (2) were interleaved with functional scans such that the functional data were acquired at approximately evenly spaced intervals throughout each imaging session.

#### 4.3.2 WF-Ca^2+^ imaging data acquisition

WF-Ca^2+^ imaging data were acquired simultaneously with all BOLD-fMRI data as we have described previously^30,34^ using CamWare software version 3.17. During each acquisition, GCaMP-sensitive (470/24nm) and GCaMP-insensitive (395/25nm) illumination (LLE 7Ch Controller from Lumencor) were interleaved at 20Hz. Exposure time for each wavelength was 40ms to avoid artifacts caused by the rolling shutter refreshing. Thus, the sequence was: 10ms blank, 40ms GCaMP-sensitive wavelength on, 10ms blank, 40ms GCaMP-insensitive wavelength on. Following data processing, where the GCaMP-insensitive channel is regressed from the GCaMP-sensitive channel, the data were at an effective 10Hz. The raw WF-Ca^2+^ imaging data had a raw spatial resolution of 25×25μm^2^ and a FOV of 1.4×1.4cm^2^ (as in ^30^).

### 4.4 Data processing

#### 4.4.1 MRI data registration and preprocessing

BOLD-fMRI data processing was performed using RABIES (Rodent automated BOLD improvement of EPI sequences^35^) v0.4.8 (https://rabies.readthedocs.io/en/stable/). All resting-state functional and isotropic MSME structural scans (from all imaging sessions) were input and processed together. Within native subject space, the BOLD-fMRI timeseries were slice time corrected, and head motion was estimated. Structural images were corrected for inhomogeneities (N3 nonparametric nonuniform intensity normalization). The within-sample anatomical template, created by averaging all structural images following non-linear registration, was registered to our in-house template (using non-linear registration), which has been previously registered to the Allen Atlas reference space CCfv3^34,36,37^.

For each BOLD-fMRI scan, a representative mean image (averaged across time) was derived. These data were corrected for intensity inhomogeneities and non-linearly registered to the corresponding isotropic structural MSME image of each mouse acquired during each imaging session. This minimized distortions caused by susceptibility artifacts. Then, the BOLD-fMRI data were moved to common space using four concatenated transformations: (1) framewise rigid head motion correction, (2) representative mean BOLD-fMRI to individual mouse/session isotropic MSME image, (3) individual mouse/session isotropic MSME image to within-dataset template, and (4) within-dataset template to out-of-sample in-house template. As part of this transformation, the data were resampled to the template resolution (0.2×0.2×0.2mm^3^). After this step, the Allen Atlas and BOLD-fMRI data resided in the same space. Registration performance was visually inspected using the RABIES quality control report. Following timeseries normalization to common space, the six-parameter motion estimates were regressed from the timeseries and used to compute framewise-displacement. Frame scrubbing for motion using a conservative 0.075mm threshold was applied. Data were filtered 0.008-0.2Hz and 30 timepoints discarded from the beginning and end of each run to avoid filter-related edge artifacts. Cerebrospinal fluid and white matter signals were regressed, and the data were smoothed with 0.4mm sigma. Global signal regression was not applied as there is no equivalent for WF-Ca^2+^ imaging data and we elected to harmonize data processing as much as possible across modalities.

#### 4.4.2 WF-Ca^2+^ imaging data registration and processing

WF-Ca^2+^ imaging data (GCaMP-sensitive and -insensitive channels) were smoothed with a 0.1mm sigma. Data were down-sampled by a factor of two in each spatial dimension and the GCaMP-insensitive channel regressed from the GCaMP-sensitive channel. ΔF/F_0_ (F = fluorescence) was computed for each run. The data were filtered, within both a slow (0.008-0.2Hz) and fast (0.4-4Hz) band. 300 timepoints were discarded from the beginning and end of each run to match the BOLD-fMRI timeseries and eliminate artifacts. Timepoints censored (based on framewise displacement) from the BOLD-fMRI timeseries were also removed from the WF-Ca^2+^ imaging timeseries. Data were registered to our in-house template using three transformations, two linear (1) WF-Ca^2+^ → Angiogram, and (2) Angiogram → MSME and one nonlinear (3) MSME to in-house template space, as described previously^30^.

#### 4.4.3 Connectomes

To compute connectomes, BOLD-fMRI or WF-Ca^2+^ imaging signals within each node (**Fig. S2**) was averaged, and the inter-regional Pearson’s correlation computed, and Fisher transformed. Connectomes were computed for each run using data residing in our in-house template space and bilateral symmetry was quantified with Spearman’s correlation.

### 4.5 Inclusion criteria

All data were temporally matched across imaging modalities. Thus, if a run (or timepoint) was excluded based on criteria applied to one modality, the corresponding simultaneously acquired data from the second modality were also excluded. Data were excluded if framewise displacement, estimated using the BOLD-fMRI timeseries, exceeded 0.075mm. If 2/3 timepoints within a run exceeded this cutoff, the run was excluded (no runs met this criterion). If registration to the in-house template space failed visual inspection, the corresponding run was excluded (6 runs, including all data from one subject). Data corruption during saving (2 instances). Low SNR, missing structural data (for data registration purposes), or other visually apparent artifacts (e.g., shutter artifacts in WF-Ca^2+^ imaging data) (12 runs). Finally, all data from a mouse were excluded if less than two runs in all sessions passed all preceding criteria (no instances). Thus, a minimum of 20mins of functional data per mouse per session were retained. Due to FOV variability, and common artifacts at the periphery of the WF-Ca^2+^ imaging data, regions at the edge of the cortical surface were excluded from WF-Ca^2+^ and cortex-only BOLD-fMRI conditions across all subjects (**Fig. S2**).

### 4.6 Analyses

#### 4.6.1 Human BOLD-fMRI data

LR resting state BOLD-fMRI data from the human connectome project S900 release were downloaded and processed following standard procedures, as described previously^18,19^. A 268-node functional atlas^5^ was used to compute connectomes using Person’s correlation and a winner-take-all ID procedure applied, as described previously^5,8^. Nodes were grouped into eight functional networks^1^.

#### 4.6.2 Connectome-based ID

The winner-take-all procedure has been described in detail previously^1^. Briefly, a query (e.g., session 1 connectomes) and target (e.g., session 2 connectomes) are selected. In an iterative process, each connectome in the query and target set are compared using Pearson’s correlation. If the highest inter-connectome correlation coefficient belongs to the same subject (i.e. within-subject correlation > all other between-subject correlations), this was denoted as a correct ID. For group ID, the mouse being queried was removed from the target session (to avoid mouse ID bleed-through). Network-based ID was performed using only nodes within a given network (**Fig. S2**). For random networks, we iteratively (N = 50) selected nodes at random and performed the ID procedure using the resulting data.

#### 4.6.3 Node and network definitions

Nodes belonging to the default mode network were selected based on the description by Whitesell and colleagues^55^. Nodes belonging to the salience and lateral cortical networks were selected based on the triple-network model^37,44,53–55^. Nodes belonging to other networks mentioned (e.g. somatosensory) were selected based on the recent comparison across species^37,53,54^.

#### 4.6.4 Edge-wise analysis

Differential power and group consistency of edges across conditions were carried out as in our previous work ^5,33^. Null distributions. To determine if the number of shared edges were greater or less than expected by chance, the number of edges identified in the real data were compared to random data where fake ‘networks’ of high consistency or differential power were generated that were symmetric and matched the size of the true results.

### 4.7 Statistics

Statistical significance tests were computed in MATLAB (R2023b & R2021bV5). Post hoc multiple comparison (MATLAB, multcompare with default “Tukey-Kramer test” to compute the p-values) was used after ANOVA test showed significance. Significant results indicated with *. Where applicable, if p values resulted lower than the arbitrary limit present within MATLAB for Anova analyses, they were set to p<0.00001, otherwise the exact p-values are reported. T-test analyses were performed with Bonferroni correction (*ttest2*, MATLAB).

## Supporting information

Supplementary Stats File

## ACKNOWLEDGEMENTS

Drs. Joel Greenwood, Omer Mano, and Paul Shamble from the Neurotechnology Core of the Yale Kavli Institute for Neuroscience for their technological expertise. Dr. Xinxin Ge for help in acquiring the multimodal imaging data. Dr. Peter Herman for performing the head-plate implant surgeries. All members of the Yale Multiscale Imaging and Spontaneous Activity in Cortex (MISAC) group for help in conceptualizing and building the in-scanner WF-Ca^2+^ imaging apparatus.

## AUTHOR CONTRIBUTIONS

FM and CH wrote the manuscript, processed the multimodal data, and conducted analyses. XS and GDG helped with analyses. WL downloaded and processed the human BOLD-fMRI data. MM processed the WF-Ca^2+^ imaging data. RTC supervised WL and multimodal data collection. XP wrote code for multimodal data processing and registration (BioImage Suite software). RFB help conceptualize and frame the project and edited the manuscript. MC edited the manuscript and supervised GDG. EMRL collected the data, conducted analyses, supervised the study, and edited the manuscript.

## DATA & CODE AVAILABILITY

Data are not available in a public repository due to ongoing work by the authors on this dataset. Access can be obtained by contacting the corresponding authors. Data from the Allen Reference Atlas and CCFv3 are available on their website: https://portal.brain-map.org/. Software packages for data processing are available: multimodal data registration and processing of human data (BioImage Suite, https://bioimagesuiteweb.github.io/webapp/), and fMRI preprocessing (RABIES, https://github.com/CoBrALab/RABIES). ID code can be found here: https://www.nitrc.org/frs/?group_id=51

