## Supplementary Stats File for "Multimodal identification of the mouse brain using simultaneous Ca^2+^ imaging and fMRI"

**Figure 3: Comparison of FC across cell-types (averaged over all animals)**

**WF-Ca^2+^_Slow_**

Anova 1 results:

| **Source** | **SS** | **df** | **MS** | **F** | **Prob>F** |
| --- | --- | --- | --- | --- | --- |
| **Groups** | 314.732483 | 4 | 78.6831206 | 672.800797 | <0.00001* |
| **Error** | 552.582201 | 4725 | 0.11694861 |  |  |
| **Total** | 867.314684 | 4729 |  |  |  |

* p-value smaller than *MATLAB* limits

Multiple comparison results:

| **Type_A** | **Type_B** | **Lower_Limit** | **A_minus_B** | **Upper_Limit** | **Pvalue** |
| --- | --- | --- | --- | --- | --- |
| GLIA | PV | 0.57086986 | 0.61376177 | 0.65665369 | 9.92E-09 |
| GLIA | SOM | 0.61859921 | 0.66149112 | 0.70438303 | 9.92E-09 |
| GLIA | VIP | 0.6479331 | 0.69082501 | 0.73371692 | 9.92E-09 |
| GLIA | SLC | 0.54577792 | 0.58866983 | 0.63156175 | 9.92E-09 |
| PV | SOM | 0.00483743 | 0.04772935 | 0.09062126 | 2.03E-02 |
| PV | VIP | 0.03417133 | 0.07706324 | 0.11995515 | 9.45E-06 |
| PV | SLC | -0.0679839 | -0.0250919 | 0.01779997 | 0.50006332 |
| SOM | VIP | -0.013558 | 0.02933389 | 0.0722258 | 3.36E-01 |
| SOM | SLC | -0.1157132 | -0.0728213 | -0.0299294 | 3.58E-05 |
| VIP | SLC | -0.1450471 | -0.1021552 | -0.0592633 | 1.07E-08 |

**WF-Ca^2+^_Fast_**

Anova 1 results:

| **Source** | **SS** | **df** | **MS** | **F** | **Prob>F** |
| --- | --- | --- | --- | --- | --- |
| **Groups** | 133.241368 | 4 | 33.3103419 | 399.201446 | 1.16E-296 |
| **Error** | 394.265519 | 4725 | 0.08344244 |  |  |
| **Total** | 527.506887 | 4729 |  |  |  |

Multiple comparison results:

| **Type_A** | **Type_B** | **Lower_Limit** | **A_minus_B** | **Upper_Limit** | **Pvalue** |
| --- | --- | --- | --- | --- | --- |
| GLIA | PV | 0.42205463 | 0.45828488 | 4.95E-01 | 9.92E-09 |
| GLIA | SOM | 0.26219368 | 0.29842393 | 3.35E-01 | 9.92E-09 |
| GLIA | VIP | 0.40446785 | 0.4406981 | 4.77E-01 | 9.92E-09 |
| GLIA | SLC | 0.34969695 | 0.3859272 | 0.42215746 | 9.92E-09 |
| PV | SOM | -0.1960912 | -0.1598609 | -1.24E-01 | 9.92E-09 |
| PV | VIP | -0.053817 | -0.0175868 | 1.86E-02 | 0.67614799 |
| PV | SLC | -0.1085879 | -0.0723577 | -3.61E-02 | 5.16E-07 |
| SOM | VIP | 0.10604392 | 0.14227417 | 1.79E-01 | 9.92E-09 |
| SOM | SLC | 0.05127302 | 0.08750327 | 1.24E-01 | 1.03E-08 |
| VIP | SLC | -0.0910011 | -0.0547709 | -1.85E-02 | 0.00035934 |

**BOLDcortex**

Anova 1 results:

| **Source** | **SS** | **df** | **MS** | **F** | **Prob>F** |
| --- | --- | --- | --- | --- | --- |
| **Groups** | 0.92078158 | 4 | 0.2301954 | 11.5800008 | 2.34E-09 |
| **Error** | 93.9268716 | 4725 | 0.0198787 |  |  |
| **Total** | 94.8476532 | 4729 |  |  |  |

Multiple comparison results:

| **Type_A** | **Type_B** | **Lower_Limit** | **A_minus_B** | **Upper_Limit** | **Pvalue** |
| --- | --- | --- | --- | --- | --- |
| GLIA | PV | -0.0341066 | -0.016423 | 0.00126067 | 0.08336796 |
| GLIA | SOM | -0.0112239 | 0.00645975 | 0.02414339 | 8.57E-01 |
| GLIA | VIP | 0.00348581 | 0.02116946 | 0.03885311 | 9.64E-03 |
| GLIA | SLC | -0.0323874 | -0.0147037 | 0.00297992 | 1.55E-01 |
| PV | SOM | 0.00519908 | 0.02288273 | 0.04056638 | 3.81E-03 |
| PV | VIP | 0.01990879 | 0.03759244 | 0.05527609 | 7.59E-08 |
| PV | SLC | -0.0159644 | 0.00171925 | 0.0194029 | 9.99E-01 |
| SOM | VIP | -0.0029739 | 0.01470972 | 0.03239337 | 0.15497857 |
| SOM | SLC | -0.0388471 | -0.0211635 | -0.0034798 | 0.00967382 |
| VIP | SLC | -0.0535568 | -0.0358732 | -0.0181895 | 3.21E-07 |

**Figure 4: FC similarity within cell-type (within cell-type ‘square’)**

**WF-Ca^2+^_Fast_**

Anova 1 results:

| **Source** | **SS** | **df** | **MS** | **F** | **Prob>F** |
| --- | --- | --- | --- | --- | --- |
| **Groups** | 27.727276 | 4 | 6.93181899 | 524.640725 | <0.00001 |
| **Error** | 57.355491 | 4341 | 0.01321251 |  |  |
| **Total** | 85.082767 | 4345 |  |  |  |

Multiple comparison results:

| **Type_A** | **Type_B** | **Lower_Limit** | **A_minus_B** | **Upper_Limit** | **Pvalue** |
| --- | --- | --- | --- | --- | --- |
| GLIA | PV | -0.1937228 | -0.1750545 | -0.1563862 | 9.92E-09 |
| GLIA | SLC | -0.2284943 | -2.09E-01 | -0.1899292 | 9.92E-09 |
| GLIA | SOM | -0.1385362 | -0.1194922 | -0.1004481 | 9.92E-09 |
| GLIA | VIP | -0.0374362 | -1.96E-02 | -0.0017912 | 0.02252669 |
| PV | SLC | -0.0492823 | -0.0341572 | -0.0190322 | 1.70E-08 |
| PV | SOM | 0.04074253 | 5.56E-02 | 0.0703821 | 9.92E-09 |
| PV | VIP | 0.14222744 | 1.55E-01 | 0.16865418 | 9.92E-09 |
| SLC | SOM | 0.07413311 | 8.97E-02 | 0.10530601 | 9.92E-09 |
| SLC | VIP | 0.1755302 | 0.18959805 | 0.20366591 | 9.92E-09 |
| SOM | VIP | 0.08613941 | 0.0998785 | 0.11361758 | 9.92E-09 |

**Figure 4: FC similarity across cell-types (cell-type ‘rows’ off diagonal)**

**WF-Ca^2+^_Fast_**

Anova 2 results:

| **Source** | **Sum Sq.** | **df** | **Mean Sq.** | **F** | **Prob>F** |
| --- | --- | --- | --- | --- | --- |
| **X1** | 43.8549057 | 4 | 1.10E+01 | 657.550877 | <0.00001 |
| **X2** | 29.6936639 | 4 | 7.42E+00 | 445.220309 | <0.00001 |
| **Error** | 350.395261 | 21015 | 1.67E-02 |  |  |
| **Total** | 423.943831 | 21023 |  |  |  |

Anova 2 with unbalanced design: X1 = session 1 and X2 = session 2

Factor X1 results

| **Type_A** | **Type_B** | **Lower_Limit** | **A_minus_B** | **Upper_Limit** | **Pvalue** |
| --- | --- | --- | --- | --- | --- |
| X1=GLIA | X1=PV | -0.1409998 | -0.132527 | -1.24E-01 | 9.92E-09 |
| X1=GLIA | X1=SLC | -0.1020095 | -0.0935944 | -8.52E-02 | 9.92E-09 |
| X1=GLIA | X1=SOM | -0.1118917 | -1.03E-01 | -9.49E-02 | 9.92E-09 |
| X1=GLIA | X1=VIP | -0.0429272 | -3.48E-02 | -0.0266679 | 9.92E-09 |
| X1=PV | X1=SLC | 0.0313413 | 3.89E-02 | 4.65E-02 | 9.92E-09 |
| X1=PV | X1=SOM | 0.02145281 | 2.91E-02 | 3.68E-02 | 9.92E-09 |
| X1=PV | X1=VIP | 0.09045577 | 9.77E-02 | 1.05E-01 | 9.92E-09 |
| X1=SLC | X1=SOM | -0.0174157 | -9.82E-03 | -2.23E-03 | 0.00380347 |
| X1=SLC | X1=VIP | 0.05159069 | 0.05879689 | 6.60E-02 | 9.92E-09 |
| X1=SOM | X1=VIP | 0.06134766 | 6.86E-02 | 7.59E-02 | 9.92E-09 |

Factor X2 results:

| **Type_A** | **Type_B** | **Lower_Limit** | **A_minus_B** | **Upper_Limit** | **Pvalue** |
| --- | --- | --- | --- | --- | --- |
| X2=GLIA | X2=PV | -0.101604 | -0.0932862 | -8.50E-02 | 9.92E-09 |
| X2=GLIA | X2=SLC | -0.1011262 | -9.24E-02 | -8.37E-02 | 9.92E-09 |
| X2=GLIA | X2=SOM | -0.1255189 | -1.17E-01 | -0.1084559 | 9.92E-09 |
| X2=GLIA | X2=VIP | -0.0580305 | -4.99E-02 | -4.18E-02 | 9.92E-09 |
| X2=PV | X2=SLC | -0.006807 | 8.90E-04 | 8.59E-03 | 9.98E-01 |
| X2=PV | X2=SOM | -0.0311722 | -2.37E-02 | -1.62E-02 | 9.92E-09 |
| X2=PV | X2=VIP | 0.03638002 | 4.34E-02 | 5.03E-02 | 9.92E-09 |
| X2=SLC | X2=SOM | -0.0325188 | -0.0245913 | -1.67E-02 | 9.92E-09 |
| X2=SLC | X2=VIP | 0.03500369 | 4.25E-02 | 4.99E-02 | 9.92E-09 |
| X2=SOM | X2=VIP | 0.05982801 | 6.71E-02 | 0.074303 | 9.92E-09 |

Anova2: Interaction of X1 and X2

| **Source** | **Sum Sq.** | **df** | **Mean Sq.** | **F** | **Prob>F** |
| --- | --- | --- | --- | --- | --- |
| **X1** | 38.765 | 4 | 9.69135 | 724.81 | <0.00001 |
| **X2** | 25.787 | 4 | 6.44681 | 482.15 | <0.00001 |
| **X1*X2** | 69.62 | 16 | 4.35124 | 325.43 | <0.00001 |
| **Error** | 280.775 | 20999 | 0.01337 |  |  |
| **Total** | 423.944 | 21023 |  |  |  |

X1 and X2 interaction results not shown

**Figure 4:** **FC similarity across cell-type (cell-type rows off diagonal)**

**WF-Ca^2+^_Fast_ and BOLDcortex**

X1 = Ca Fast and X2 = BOLD

| **Source** | **Sum Sq.** | **df** | **Mean Sq.** | **F** | **Prob>F** |
| --- | --- | --- | --- | --- | --- |
| **X1** | 9.095 | 4 | 2.2737 | 228.98 | <0.00001 |
| **X2** | 4.64E+01 | 4 | 11.6119 | 1.17E+03 | <0.00001 |
| **Error** | 2.12E+02 | 2.13E+04 | 9.90E-03 |  |  |
| **Total** | 2.67E+02 | 2.13E+04 |  |  |  |

Factor X1 results:

| **Session_I** | **Session_II** | **Lower_Limit** | **A_minus_B** | **Upper_Limit** | **Pvalue** |
| --- | --- | --- | --- | --- | --- |
| X1=GLIA | X1=PV | 0.01563957 | 2.20E-02 | 0.02838916 | 9.92E-09 |
| X1=GLIA | X1=SLC | -4.38E-02 | -0.0370883 | -0.0303975 | 9.92E-09 |
| X1=GLIA | X1=SOM | -2.25E-02 | -0.0159789 | -0.0094403 | 1.02E-08 |
| X1=GLIA | X1=VIP | -2.92E-02 | -0.0230269 | -0.0168124 | 9.92E-09 |
| X1=PV | X1=SLC | -0.0650017 | -0.0591026 | -0.0532036 | 9.92E-09 |
| X1=PV | X1=SOM | -0.0437191 | -0.0379933 | -0.0322674 | 9.92E-09 |
| X1=PV | X1=VIP | -0.0503941 | -0.0450413 | -0.0396885 | 9.92E-09 |
| X1=SLC | X1=SOM | 0.01503365 | 0.02110935 | 0.02718504 | 9.92E-09 |
| X1=SLC | X1=VIP | 0.00833585 | 0.01406133 | 0.01978681 | 1.01E-08 |
| X1=SOM | X1=VIP | -0.0125949 | -0.007048 | -0.0015012 | 4.80E-03 |

Factor X2 results:

| **Session_I** | **Session_II** | **Lower_Limit** | **A_minus_B** | **Upper_Limit** | **Pvalue** |
| --- | --- | --- | --- | --- | --- |
| X2=GLIA | X2=PV | -0.1298145 | -0.1234397 | -0.1170649 | 9.92E-09 |
| X2=GLIA | X2=SLC | -0.1529028 | -0.146212 | -0.1395212 | 9.92E-09 |
| X2=GLIA | X2=SOM | -0.0980439 | -0.0915053 | -0.0849667 | 9.92E-09 |
| X2=GLIA | X2=VIP | -0.0645592 | -0.0583447 | -0.0521302 | 9.92E-09 |
| X2=PV | X2=SLC | -0.0286714 | -0.0227724 | -0.0168733 | 9.92E-09 |
| X2=PV | X2=SOM | 0.02620856 | 0.03193441 | 0.03766026 | 9.92E-09 |
| X2=PV | X2=VIP | 0.05974216 | 0.06509495 | 0.07044775 | 9.92E-09 |
| X2=SLC | X2=SOM | 0.04863109 | 5.47E-02 | 0.06078248 | 9.92E-09 |
| X2=SLC | X2=VIP | 8.21E-02 | 8.79E-02 | 0.0935928 | 9.92E-09 |
| X2=SOM | X2=VIP | 0.02761368 | 3.32E-02 | 0.0387074 | 9.92E-09 |

Anova2 for interaction:

| **Source** | **Sum Sq.** | **df** | **Mean Sq.** | **F** | **Prob>F** |
| --- | --- | --- | --- | --- | --- |
| **X1** | 9.01256313 | 4 | 2.25314078 | 2.27E+02 | 1.86E-191 |
| **X2** | 44.4140483 | 4 | 11.1035121 | 1.12E+03 | 0.00E+00 |
| **X1*X2** | 0.56757526 | 16 | 3.55E-02 | 3.58E+00 | 1.53E-06 |
| **Error** | 211.00476 | 21291 | 9.91E-03 |  |  |
| **Total** | 267.114866 | 21315 |  |  |  |

X1 and X2 interaction results not shown

**Figure 5: ID rate of mouse (between pairs of sessions) - session effect**

**WF-Ca^2+^_Slow_**

Anova2

| **Source** | **Sum Sq.** | **df** | **Mean Sq.** | **F** | **Prob>F** |
| --- | --- | --- | --- | --- | --- |
| **X1** | 5619.8409 | 2 | 2.81E+03 | 8.88E+02 | 1.39E-125 |
| **X2** | 7937.16539 | 2 | 3968.5827 | 1.25E+03 | 5.58E-145 |
| **Error** | 933.152304 | 295 | 3.16E+00 |  |  |
| **Total** | 10207.2397 | 299 |  |  |  |

Anova 2: X1 = session 1—reference; X2 = session 2—target

X1 results:

| **X1** | **X1** | **Lower_Limit** | **Difference** | **Upper_Limit** | **Pvalue** |
| --- | --- | --- | --- | --- | --- |
| X1=s1 | X1=s3 | -7.4440313 | -6.7633375 | -6.0826438 | 9.56E-10 |
| X1=s1 | X1=s2 | -12.899204 | -12.21851 | -11.537817 | 9.56E-10 |
| X1=s3 | X1=s2 | -6.1358664 | -5.4551727 | -4.774479 | 9.56E-10 |

X2 results:

| **X2** | **X2** | **Lower_Limit** | **Difference** | **Upper_Limit** | **Pvalue** |
| --- | --- | --- | --- | --- | --- |
| X2=s3 | X2=s1 | 9.10364603 | 9.78433976 | 10.4650335 | 9.56E-10 |
| X2=s3 | X2=s2 | -5.1128645 | -4.4321708 | -3.7514771 | 9.56E-10 |
| X2=s1 | X2=s2 | -14.897204 | -14.216511 | -13.535817 | 9.56E-10 |

Anova for interaction

| **Source** | **Sum Sq.** | **df** | **Mean Sq.** | **F** | **Prob>F** |
| --- | --- | --- | --- | --- | --- |
| **Reference** | 0 | 0 | 0 | 0 | NaN |
| **Target** | 0 | 0 | 0 | 0 | NaN |
| **Reference*Target** | 86.8 | 1.00E+00 | 86.8342 | 30.17 | 8.59E-08 |
| **Error** | 846.3 | 2.94E+02 | 2.8786 |  |  |
| **Total** | 10207.2 | 2.99E+02 |  |  |  |

X1 and X2 interaction:

| **Combination 1** | **Combination 2** | **Lower_Limit** | **Difference** | **Upper_Limit** | **Pvalue** |
| --- | --- | --- | --- | --- | --- |
| Reference=s1,Target=s3 | Reference=s2,Target=s3 | -12.195016 | -11.142504 | -10.089992 | 8.97E-08 |
| Reference=s1,Target=s3 | Reference=s3,Target=s1 | 3.04449636 | 4.09700867 | 5.14952097 | 8.97E-08 |
| Reference=s1,Target=s3 | Reference=s2,Target=s1 | -3.4866828 | -2.4341705 | -1.3816582 | 8.97E-08 |
| Reference=s1,Target=s3 | Reference=s1,Target=s2 | -4.4086767 | -3.3561644 | -2.3036521 | 8.97E-08 |
| Reference=s1,Target=s3 | Reference=s3,Target=s2 | -12.248021 | -11.195508 | -10.142996 | 8.97E-08 |
| Reference=s2,Target=s3 | Reference=s3,Target=s1 | 14.1870002 | 15.2395125 | 16.2920248 | 8.97E-08 |
| Reference=s2,Target=s3 | Reference=s2,Target=s1 | 7.65582103 | 8.70833333 | 9.76084564 | 8.97E-08 |
| Reference=s2,Target=s3 | Reference=s1,Target=s2 | 6.73382712 | 7.78633942 | 8.83885172 | 8.97E-08 |
| Reference=s2,Target=s3 | Reference=s3,Target=s2 | -1.1055168 | -0.0530045 | 0.99950777 | 0.99999999 |
| Reference=s3,Target=s1 | Reference=s2,Target=s1 | -7.5836914 | -6.5311791 | -5.4786668 | 8.97E-08 |
| Reference=s3,Target=s1 | Reference=s1,Target=s2 | -8.5056854 | -7.4531731 | -6.4006607 | 8.97E-08 |
| Reference=s3,Target=s1 | Reference=s3,Target=s2 | -16.345029 | -15.292517 | -14.240005 | 8.97E-08 |
| Reference=s2,Target=s1 | Reference=s1,Target=s2 | -1.9745062 | -0.9219939 | 0.13051839 | 0.14163948 |
| Reference=s2,Target=s1 | Reference=s3,Target=s2 | -9.8138502 | -8.7613379 | -7.7088256 | 8.97E-08 |
| Reference=s1,Target=s2 | Reference=s3,Target=s2 | -8.8918563 | -7.839344 | -6.7868317 | 8.97E-08 |

**Figure 5: ID rate of cell-type (between pairs of sessions) - session effect**

**WF-Ca^2+^_Slow_**

X1 session 1; X2 session 2

| **Source** | **Sum Sq.** | **df** | **Mean Sq.** | **F** | **Prob>F** |
| --- | --- | --- | --- | --- | --- |
| **X1** | 2918.28259 | 2 | 1459.14129 | 231.603743 | 3.39E-61 |
| **X2** | 961.924522 | 2 | 480.962261 | 76.3412427 | 1.91E-27 |
| **Error** | 1858.54804 | 295 | 6.30016284 |  |  |
| **Total** | 5027.26397 | 299 |  |  |  |

X1 results:

| **X1** | **X1** | **Lower_Limit** | **Difference** | **Upper_Limit** | **Pvalue** |
| --- | --- | --- | --- | --- | --- |
| X1=s1 | X1=s3 | -4.72E+00 | -3.7601044 | -2.7994606 | 9.56E-10 |
| X1=s1 | X1=s2 | -9.75E+00 | -8.791048 | -7.8304042 | 9.56E-10 |
| X1=s3 | X1=s2 | -5.9915874 | -5.0309436 | -4.0702998 | 9.56E-10 |

X2 results:

| **X2** | **X2** | **Lower_Limit** | **Difference** | **Upper_Limit** | **Pvalue** |
| --- | --- | --- | --- | --- | --- |
| X2=s3 | X2=s1 | 2.99764165 | 3.95828545 | 4.91892924 | 9.56E-10 |
| X2=s3 | X2=s2 | -1.7178257 | -0.7571819 | 0.20346185 | 0.1543611 |
| X2=s1 | X2=s2 | -5.6761112 | -4.7154674 | -3.7548236 | 9.56E-10 |

Anova for interaction

| **Source** | **Sum Sq.** | **df** | **Mean Sq.** | **F** | **Prob>F** |
| --- | --- | --- | --- | --- | --- |
| **Reference** | 0 | 0 | 0 | 0 | NaN |
| **Target** | 0 | 0 | 0 | 0 | NaN |
| **Reference*Target** | 389.1 | 1 | 389.104 | 77.85 | 1.01E-16 |
| **Error** | 1469.44 | 294 | 4.998 |  |  |
| **Total** | 5027.26 | 299 |  |  |  |

X1 and X2 interaction:

| **Combination 1** | **Combination 2** | **Lower_Limit** | **Difference** | **Upper_Limit** | **Pvalue** |
| --- | --- | --- | --- | --- | --- |
| Reference=s1,Target=s3 | Reference=s2,Target=s3 | -7.9001918 | -6.5133181 | -5.13E+00 | 8.97E-08 |
| Reference=s1,Target=s3 | Reference=s3,Target=s1 | 1.08903726 | 2.47591091 | 3.86E+00 | 1.17E-06 |
| Reference=s1,Target=s3 | Reference=s2,Target=s1 | -6.2196362 | -4.8327626 | -3.45E+00 | 8.97E-08 |
| Reference=s1,Target=s3 | Reference=s1,Target=s2 | 0.13367429 | 1.52054795 | 2.91E+00 | 0.01936466 |
| Reference=s1,Target=s3 | Reference=s3,Target=s2 | -5.90416 | -4.5172864 | -3.1304127 | 8.97E-08 |
| Reference=s2,Target=s3 | Reference=s3,Target=s1 | 7.60235537 | 8.98922902 | 10.3761027 | 8.97E-08 |
| Reference=s2,Target=s3 | Reference=s2,Target=s1 | 0.2936819 | 1.68055556 | 3.06742921 | 0.00536393 |
| Reference=s2,Target=s3 | Reference=s1,Target=s2 | 6.6469924 | 8.03386606 | 9.42073971 | 8.97E-08 |
| Reference=s2,Target=s3 | Reference=s3,Target=s2 | 0.60915809 | 1.99603175 | 3.3829054 | 0.00027606 |
| Reference=s3,Target=s1 | Reference=s2,Target=s1 | -8.6955471 | -7.3086735 | -5.9217998 | 8.97E-08 |
| Reference=s3,Target=s1 | Reference=s1,Target=s2 | -2.3422366 | -0.955363 | 0.43151069 | 0.44801233 |
| Reference=s3,Target=s1 | Reference=s3,Target=s2 | -8.3800709 | -6.9931973 | -5.6063236 | 8.97E-08 |
| Reference=s2,Target=s1 | Reference=s1,Target=s2 | 4.96643685 | 6.3533105 | 7.74018416 | 8.97E-08 |
| Reference=s2,Target=s1 | Reference=s3,Target=s2 | -1.0713975 | 0.31547619 | 1.70234985 | 0.99873097 |
| Reference=s1,Target=s2 | Reference=s3,Target=s2 | -7.424708 | -6.0378343 | -4.6509607 | 8.97E-08 |

**Figure 5: ID rate of mouse (between pairs of sessions) - session effect**

**WF-Ca^2+^_Fast_**

Anova2: X1 = session 1; X2 = session 2

| **Source** | **Sum Sq.** | **df** | **Mean Sq.** | **F** | **Prob>F** |
| --- | --- | --- | --- | --- | --- |
| **X1** | 4412.0286 | 2 | 2.21E+03 | 813.026974 | 9.48E-121 |
| **X2** | 9192.47447 | 2 | 4.60E+03 | 1693.94408 | 1.93E-162 |
| **Error** | 800.433736 | 295 | 2.7133347 |  |  |
| **Total** | 10956.9378 | 299 |  |  |  |

X1 results:

| **X1** | **X1** | **Lower_Limit** | **Difference** | **Upper_Limit** | **Pvalue** |
| --- | --- | --- | --- | --- | --- |
| X1=s1 | X1=s3 | -1.15E+01 | -1.08E+01 | -10.21527 | 9.56E-10 |
| X1=s1 | X1=s2 | -6.19E+00 | -5.56E+00 | -4.9289547 | 9.56E-10 |
| X1=s3 | X1=s2 | 4.66E+00 | 5.29E+00 | 5.91674673 | 9.56E-10 |

X2 results:

| **X2** | **X2** | **Lower_Limit** | **Difference** | **Upper_Limit** | **Pvalue** |
| --- | --- | --- | --- | --- | --- |
| X2=s3 | X2=s1 | 14.0096244 | 14.6400563 | 15.2704882 | 9.56E-10 |
| X2=s3 | X2=s2 | 1.88324085 | 2.51367274 | 3.14410464 | 9.56E-10 |
| X2=s1 | X2=s2 | -12.756815 | -12.126384 | -11.495952 | 9.56E-10 |

Anova: Interactions

| **Source** | **Sum Sq.** | **df** | **Mean Sq.** | **F** | **Prob>F** |
| --- | --- | --- | --- | --- | --- |
| **Reference** | 0 | 0 | 0 | 0 | NaN |
| **Target** | 0 | 0 | 0 | 0 | NaN |
| **Reference*Target** | 13.1 | 1 | 13.0939 | 4.89 | 0.0278 |
| **Error** | 787.3 | 294 | 2.678 |  |  |
| **Total** | 10956.9 | 299 |  |  |  |

X1 and X2 interaction:

| **Combination 1** | **Combination 2** | **Lower_Limit** | **Difference** | **Upper_Limit** | **Pvalue** |
| --- | --- | --- | --- | --- | --- |
| Reference=s1,Target=s3 | Reference=s2,Target=s3 | -6.16E+00 | -5.14E+00 | -4.1263762 | 8.97E-08 |
| Reference=s1,Target=s3 | Reference=s3,Target=s1 | 3.20E+00 | 4.21218899 | 5.22736532 | 8.97E-08 |
| Reference=s1,Target=s3 | Reference=s2,Target=s1 | 8.06549337 | 9.08066971 | 10.0958461 | 8.97E-08 |
| Reference=s1,Target=s3 | Reference=s1,Target=s2 | 1.91633051 | 2.93150685 | 3.94668319 | 8.97E-08 |
| Reference=s1,Target=s3 | Reference=s3,Target=s2 | -9.347205 | -8.3320287 | -7.3168524 | 8.97E-08 |
| Reference=s2,Target=s3 | Reference=s3,Target=s1 | 8.33856516 | 9.3537415 | 10.3689178 | 8.97E-08 |
| Reference=s2,Target=s3 | Reference=s2,Target=s1 | 13.2070459 | 14.2222222 | 1.52E+01 | 8.97E-08 |
| Reference=s2,Target=s3 | Reference=s1,Target=s2 | 7.05788302 | 8.07305936 | 9.0882357 | 8.97E-08 |
| Reference=s2,Target=s3 | Reference=s3,Target=s2 | -4.2056525 | -3.1904762 | -2.1752999 | 8.97E-08 |
| Reference=s3,Target=s1 | Reference=s2,Target=s1 | 3.85330439 | 4.86848073 | 5.88365706 | 8.97E-08 |
| Reference=s3,Target=s1 | Reference=s1,Target=s2 | -2.2958585 | -1.2806821 | -0.2655058 | 0.00294027 |
| Reference=s3,Target=s1 | Reference=s3,Target=s2 | -13.559394 | -12.544218 | -11.529041 | 8.97E-08 |
| Reference=s2,Target=s1 | Reference=s1,Target=s2 | -7.1643392 | -6.1491629 | -5.1339865 | 8.97E-08 |
| Reference=s2,Target=s1 | Reference=s3,Target=s2 | -18.427875 | -17.412698 | -16.397522 | 8.97E-08 |
| Reference=s1,Target=s2 | Reference=s3,Target=s2 | -12.278712 | -11.263536 | -10.248359 | 8.97E-08 |

**Figure 5: ID rate of cell-type (between pairs of sessions) - session effect**

**WF-Ca^2+^_Fast_**

Anova2: X1 = session 1; X2 = session 2

| **Source** | **Sum Sq.** | **df** | **Mean Sq.** | **F** | **Prob>F** |
| --- | --- | --- | --- | --- | --- |
| **X1** | 1702.64898 | 2 | 851.32449 | 128.536749 | 7.15E-41 |
| **X2** | 1755.32633 | 2 | 877.663163 | 132.513479 | 8.67E-42 |
| **Error** | 1953.84376 | 295 | 6.62319919 |  |  |
| **Total** | 4264.96515 | 299 |  |  |  |

X1 results:

| **X1** | **X1** | **Lower_Limit** | **Difference** | **Upper_Limit** | **Pvalue** |
| --- | --- | --- | --- | --- | --- |
| X1=s1 | X1=s3 | -7.65E+00 | -6.663299 | -5.6783349 | 9.56E-10 |
| X1=s1 | X1=s2 | -5.1845507 | -4.1995866 | -3.2146225 | 9.56E-10 |
| X1=s3 | X1=s2 | 1.47874825 | 2.46371235 | 3.44867645 | 1.46E-08 |

X2 results:

| **X2** | **X2** | **Lower_Limit** | **Difference** | **Upper_Limit** | **Pvalue** |
| --- | --- | --- | --- | --- | --- |
| X2=s3 | X2=s1 | 5.6938843 | 6.6788484 | 7.66381251 | 9.56E-10 |
| X2=s3 | X2=s2 | 1.06944472 | 2.05E+00 | 3.03937293 | 3.04E-06 |
| X2=s1 | X2=s2 | -5.6094037 | -4.6244396 | -3.6394755 | 9.56E-10 |

Anova for interaction

| **Source** | **Sum Sq.** | **df** | **Mean Sq.** | **F** | **Prob>F** |
| --- | --- | --- | --- | --- | --- |
| **Reference** | 0 | 0 | 0 | 0 | NaN |
| **Target** | 0 | 0 | 0 | 0 | NaN |
| **Reference*Target** | 1292.35 | 1 | 1292.35 | 574.39 | 4.10E-71 |
| **Error** | 661.49 | 294 | 2.25 |  |  |
| **Total** | 4264.97 | 299 |  |  |  |

X1 and X2 interaction:

| **Combination 1** | **Combination 2** | **Lower_Limit** | **Difference** | **Upper_Limit** | **Pvalue** |
| --- | --- | --- | --- | --- | --- |
| Reference=s1,Target=s3 | Reference=s2,Target=s3 | -0.9790279 | -4.85E-02 | 8.82E-01 | 1.00E+00 |
| Reference=s1,Target=s3 | Reference=s3,Target=s1 | 3.2361082 | 4.17E+00 | 5.10E+00 | 8.97E-08 |
| Reference=s1,Target=s3 | Reference=s2,Target=s1 | 1.54874992 | 2.48E+00 | 3.41E+00 | 8.97E-08 |
| Reference=s1,Target=s3 | Reference=s1,Target=s2 | 5.27496758 | 6.21E+00 | 7.13599133 | 8.97E-08 |
| Reference=s1,Target=s3 | Reference=s3,Target=s2 | -5.539402 | -4.6088901 | -3.6783783 | 8.97E-08 |
| Reference=s2,Target=s3 | Reference=s3,Target=s1 | 3.28462418 | 4.21513605 | 5.14564793 | 8.97E-08 |
| Reference=s2,Target=s3 | Reference=s2,Target=s1 | 1.5972659 | 2.52777778 | 3.45828965 | 8.97E-08 |
| Reference=s2,Target=s3 | Reference=s1,Target=s2 | 5.32348356 | 6.25399543 | 7.18450731 | 8.97E-08 |
| Reference=s2,Target=s3 | Reference=s3,Target=s2 | -5.49E+00 | -4.5603741 | -3.6298623 | 8.97E-08 |
| Reference=s3,Target=s1 | Reference=s2,Target=s1 | -2.6178702 | -1.6873583 | -0.7568464 | 7.44E-07 |
| Reference=s3,Target=s1 | Reference=s1,Target=s2 | 1.1083475 | 2.03885938 | 2.96937126 | 9.00E-08 |
| Reference=s3,Target=s1 | Reference=s3,Target=s2 | -9.7060221 | -8.7755102 | -7.8449983 | 8.97E-08 |
| Reference=s2,Target=s1 | Reference=s1,Target=s2 | 2.79570578 | 3.72621766 | 4.65672953 | 8.97E-08 |
| Reference=s2,Target=s1 | Reference=s3,Target=s2 | -8.0186638 | -7.0881519 | -6.1576401 | 8.97E-08 |
| Reference=s1,Target=s2 | Reference=s3,Target=s2 | -11.744881 | -10.81437 | -9.8838577 | 8.97E-08 |

**Figure 5: ID rate of mouse across conditions**

Comparing ID rate of animals using WF-Ca^2+^_Slow_, WF-Ca^2+^_Fast_, and BOLDcortex

Anova 1

| **Source** | **SS** | **df** | **MS** | **F** | **Prob>F** |
| --- | --- | --- | --- | --- | --- |
| **Groups** | 32105.8871 | 2 | 16052.9435 | 622.01523 | 3.49E-170 |
| **Error** | 23149.7392 | 897 | 25.807959 |  |  |
| **Total** | 55255.6263 | 899 |  |  |  |

X1 has three levels: WF-Ca^2+^_Slow_, WF-Ca^2+^_Fast_, and BOLDcortex

Multiple comparison:

X1 results:

| **Group_I** | **Group_II** | **Lower_Limit** | **Difference** | **Upper_Limit** | **Pvalue** |
| --- | --- | --- | --- | --- | --- |
| low_40 | high_40 | -7.4399698 | -6.47E+00 | -5.4956696 | 9.56E-10 |
| low_40 | bold_40 | 7.15857448 | 8.13E+00 | 9.1028747 | 9.56E-10 |
| high_40 | bold_40 | 13.6263942 | 1.46E+01 | 15.5706944 | 9.56E-10 |

**Figure 5: ID rate of cell-type across conditions**

Comparing ID rate of animals using WF-Ca^2+^_Slow_, WF-Ca^2+^_Fast_, and BOLDcortex

Anova 1

| **Source** | **SS** | **df** | **MS** | **F** | **Prob>F** |
| --- | --- | --- | --- | --- | --- |
| **Groups** | 196784.53 | 2 | 98392.2652 | 6046.46824 | <0.00001 |
| **Error** | 14596.5973 | 897 | 16.2726837 |  |  |
| **Total** | 211381.128 | 899 |  |  |  |

Multiple comparisons

X1 results:

| **Group_I** | **Group_II** | **Lower_Limit** | **Difference** | **Upper_Limit** | **Pvalue** |
| --- | --- | --- | --- | --- | --- |
| cell_low_40 | cell_high_40 | -13.962714 | -13.190769 | -12.418825 | 9.56E-10 |
| cell_low_40 | cell_bold_40 | 21.8461156 | 22.6180601 | 23.3900046 | 9.56E-10 |
| cell_high_40 | cell_bold_40 | 35.0368849 | 35.8088294 | 36.580774 | 9.56E-10 |
