## Supplementary Material for "Multimodal identification of the mouse brain using simultaneous Ca^2+^ imaging and fMRI"

### SUPPLEMENTARY METHODS:

#### Terminology

- **Group**: Each of the five genotypes of mice (SLC, VIP, PV, SOM, GLIA).
- **Condition**: Each of the four types of signal analyzed (WF-$Ca^{2+}_{Slow}$, WF-$Ca^{2+}_{Fast}$, BOLD-fMRI cortex and BOLD-fMRI whole-brain.
- **Inter-condition**: Interaction across two or more or the conditions.
- **Intra-condition**: Comparisons within the same experimental condition (usually across sessions).
- **Inter-connectome**: Interactions between distinct functional connectomes.
- **Session:** One of three imaging timepoints. Imaging days were spaced a minimum of 7-days apart. From each session, a minimum of 2 ‘runs’ per mouse (and maximum of 5) were retained for our analyses (see inclusion criteria).
- **Run:** Ten-minutes of functional data collected during one scan (see imaging protocol).

### SUPPLEMENTARY FIGURES

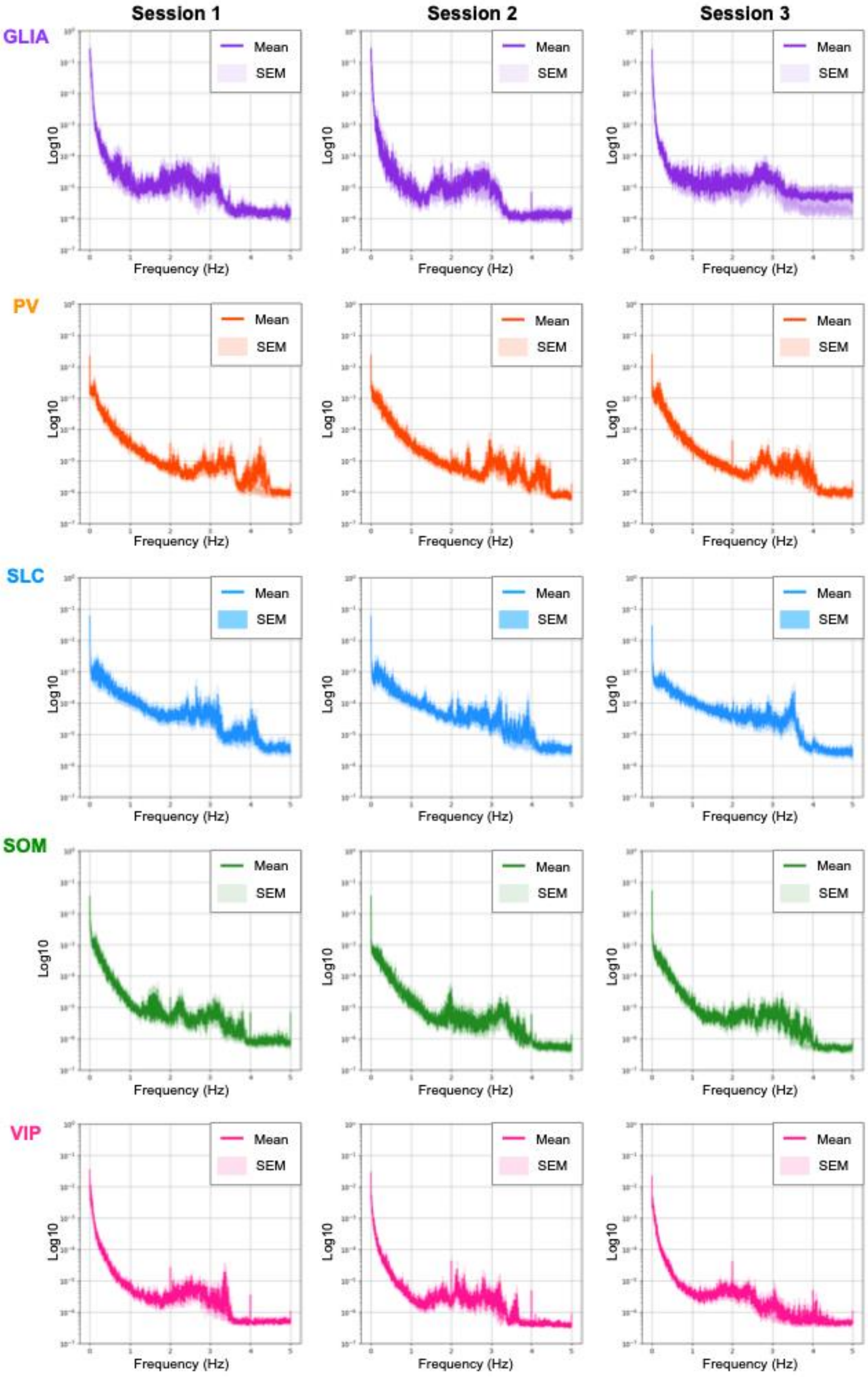


***Fig. S1 | Unfiltered WF-Ca$^{2+}$ imaging data power spectra by session and group.*** *The mean power spectrum for each group was computed across all runs, for each session. Data are displayed with +/- 1 standard error of the mean (SEM) as shading.*

| ## | Nodes Allen CCfv3 | 2D | 3D |
|---|---|---|---|
| 1 | Frontal pole, cerebral cortex (FRP,184) | | 1 |
| 2 | Primary motor area (MOp,985) | 1,4,6 | 1,6,7,9 |
| 3 | Secondary motor area (MOs,993) | 1,4,6 | 1,2,6,7,9 |
| 4 | Primary somatosensory area, nose (SSp-n,353) | 1,2,4,6 | 1,2,6,7,9 |
| 5 | Primary somatosensory area, barrel field (S. Sp-bfd,329) | 1,2,4,6 | 1,2,6,7,9 |
| 6 | Primary somatosensory area, lower limb (SSp-ll,337) | 1,2,4,6 | 1,2,6,7,9 |
| 7 | Primary somatosensory area, mouth (SSp-m,345) | 1,2,4,6 | 1,2,6,7,9 |
| 8 | Primary somatosensory area, upper limb (SSp-ul,369) | 1,2,4,6 | 1,2,6,7,9 |
| 9 | Primary somatosensory area, trunk (SSp-tr,361) | 1,2,4,6 | 1,2,6,7,9 |
| 10 | Primary somatosensory area, unassigned (SSp-un,182305689) | 1,2,4,6 | 1,2,6,7,9 |
| 11 | Supplemental somatosensory area (SSs,378) | 1,2,4,6 | 1,2,6,7,9 |
| 12 | Gustatory areas (GU,1057) | | 1 |
| 13 | Visceral area (VISC,677) | | 1 |
| 14 | Dorsal auditory area (AUDd,1011) | 1,4 | 1,3,6,7 |
| 15 | Primary auditory area (AUDp,1002) | | 1,3,6,7 |
| 16 | Posterior auditory area (AUDpo,1027) | | 1,3,6,7 |
| 17 | Ventral auditory area (AUDv,1018) | | 1,3,6,7 |
| 18 | Anterolateral visual area (VISal,402) | 1,3,4 | 1,4,6,7 |
| 19 | Anteromedial visual area (VISam,394) | 1,3,4 | 1,4,6,7 |
| 20 | Lateral visual area (VISl,409) | 1,3,4 | 1,4,6,7 |
| 21 | Primary visual area (VISp,385) | 1,3,4 | 1,4,6,7 |
| 22 | Posterolateral visual area (VISpl,425) | | 1,4,6,7 |
| 23 | Posteromedial visual area (VISpm,533) | 1,3,4 | 1,4,6,7 |
| 24 | Anterior area (VISa,312782546) | 1,3,4 | 1,4,6,7 |
| 25 | Laterointermediate area (VISli,312782574) | | 1,4,6,7 |
| 26 | Rostrolateral visual area (VISrl,417) | 1,3,4 | 1,4,6,7 |
| 27 | Postrhinal area (VISpor,312782628) | | 1,4,7 |
| 28 | Anterior cingulate area, dorsal part (ACAd,39) | 1,5,7 | 1,7,8,10,11 |
| 29 | Anterior cingulate area, ventral part (ACAv,48) | | 1,8,10,11 |
| 30 | Prelimbic area (PL,972) | | 1,7,10 |
| 31 | Infralimbic area (ILA,44) | | 1,10 |
| 32 | Orbital area, lateral part (ORBl,723) | | 1 |
| 33 | Orbital area, medial part (ORBm,731) | | 1 |
| 34 | Orbital area, ventrolateral part (ORBvl,746) | | 1 |
| 35 | Agranular insular area, dorsal part (AId,104) | | 1,11 |
| 36 | Agranular insular area, posterior part (AIp,111) | | 1,11 |
| 37 | Agranular insular area, ventral part (AIv,119) | | 1,11 |
| 38 | Retrosplenial area, lateral agranular part (RSPagl,894) | 1,3,4,5,7 | 1,4,6,7,8,10 |
| 39 | Retrosplenial area, dorsal part (RSPd,879) | 1,5,7 | 1,7,8,10 |
| 40 | Retrosplenial area, ventral part (RSPv,886) | 1,5,7 | 1,7,8,10 |
| 41 | Temporal association areas (TEa,541) | | 1,7,9,10 |
| 42 | Perirhinal area (PERI,922) | | 1,8,10 |
| 43 | Ectorhinal area (ECT,895) | Cortical | 1,6,8,10 |
| 44 | Main olfactory bulb (MOB,507) | Subcortical | 1,5,6 |
| 45 | Accessory olfactory bulb (AOB,151) | | 1,5,6 |
| 46 | Anterior olfactory nucleus (AON,159) | | 1,5,6 |
| 47 | Taenia tecta (TT,589) | | 1,5,6 |
| 48 | Dorsal peduncular area (DP,814) | | 1,5,6 |
| 49 | Piriform area (PIR,961) | | 1,5,6 |
| 50 | Nucleus of the lateral olfactory tract (NLOT,619) | | 1,5,6 |
| 51 | Cortical amygdalar area (COA,631) | | 1,8 |
| 52 | Piriform-amygdalar area (PAA,788) | | 1,8 |
| 53 | Postpiriform transition area (TR,566) | | 1,5,6 |
| 54 | Hippocampal region (HIP,1080) | | 1,8,10 |
| 55 | Retrohippocampal region (RHP,822) | | 1,8,10 |
| 56 | Claustrum (CLA,583) | | 1 |
| 57 | Endopiriform nucleus (EP,942) | | 1 |
| 58 | Lateral amygdalar nucleus (LA,131) | | 1,8 |
| 59 | Basolateral amygdalar nucleus (BLA,295) | | 1,8 |
| 60 | Basomedial amygdalar nucleus (BMA,319) | | 1,8 |
| 61 | Posterior amygdalar nucleus (PA,780) | | 1,8 |
| 62 | Striatum dorsal region (STRd,485) | | 1,10 |
| 63 | Striatum ventral region (STRv,493) | | 1,11 |
| 64 | Lateral septal complex (LSX,275) | | 1,11 |
| 65 | Striatum-like amygdalar nuclei (sAMY,278) | | 1,11 |

| # | Nodes Allen CCfv3 | 3D networks |
|---|---|---|
| 66 | Pallidum, dorsal region (PALd,818) | 1 |
| 67 | Pallidum, ventral region (PALv,835) | 1 |
| 68 | Pallidum, medial region (PALm,826) | 1 |
| 69 | Pallidum, caudal region (PALc,809) | 1 |
| 70 | Thalamus, sensory-motor cortex (DORsm,864) | 1,10 |
| 71 | Thalamus, polymodal (DORpm,856) | 1,10 |
| 72 | Periventricular zone (PVZ,157) | 1,12,13 |
| 73 | Periventricular region (PVR,141) | 1,12,13 |
| 74 | Hypothalamic medial zone (MEZ,467) | 1,8,11 |
| 75 | Hypothalamic lateral zone (LZ,290) | 1,8,11 |
| 76 | Midbrain, sensory related (MBsen,339) | 1,12,13 |
| 77 | Midbrain, motor related (MBmot,323) | 1,12,13 |
| 78 | Midbrain, behavioral state related (MBsta,348) | 1,12,13 |
| 79 | Pons, sensory related (P-sen,1132) | 1,12,13 |
| 80 | Pons, motor related (P-mot,987) | 1,12,13 |
| 81 | Pons, behavioral state related (P-sat,1117) | 1,12,13 |
| 82 | Medulla, sensory related (MY-sen,386) | 1,12,13 |
| 83 | Medulla, motor related (MY-mot,370) | 1,12,13 |
| 84 | Medulla, behavioral state related (MY-sat,379) | 1,12,13 |
| 85 | Cerebellar cortex (CBX,528) | 1,12 |
| 86 | Cerebellar nuclei (CBN,519) | 1,12 |

**2D networks**
1.All nodes
2.Somatosensory
3.Visual
4.Sensory (all)
5.Limbic
6.LCN
7.DMN

**3D networks**
1.All nodes
2.Somatosensory
3.Auditory
4.Visual
5.Olfactory
6.Sensory (all)
7.Cortex
8.Limbic
9.LCN
10.DMN
11.Salience
12.Subcortex
13.Braintem

Matched between modalities
Excluded due to imperfect coverage

**Network color-coding**

| | |
|---|---|
| Somatosensory | Limbic |
| Sensory auditory | Lateral cortical |
| Sensory visual | Default mode network |
| Sensory olfactory | Salience |
| Sensory (all) | Subcortex |
| Cortex | Brainstem |

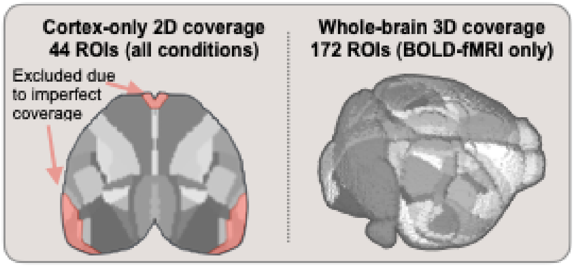


*Fig. S2 | **Nodes from Allen Institute used for cortex-only and whole-brain connectomes and canonical networks.** Nodes are listed with cortical regions above the line (1-44) and subcortical nodes below the line (44-86). Membership in 2D and 3D canonical networks indicated by number and color. Overlap across modalities as indicated by boxed nodes. Nodes with obstructions in WF-$Ca^{2+}$ data appear in red and are excluded (grey boxed inlay).*

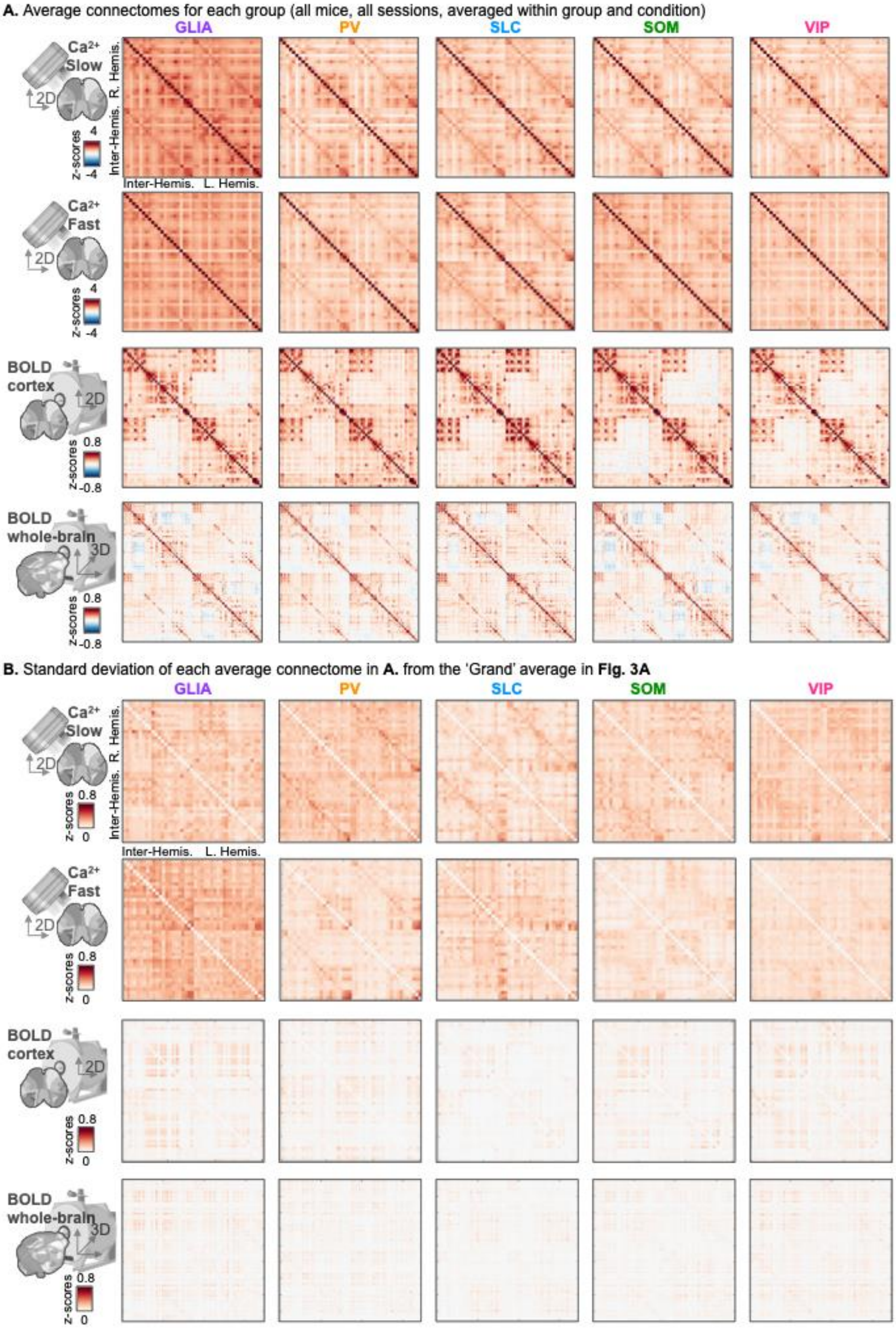


***Fig. S3 | Average connectomes within groups and their standard deviation from the 'grand' average connectomes.*** ***A.*** *Average connectomes obtained in each group (left to right: GLIA, PV, SLC, SOM, VIP), for each condition (top to bottom: WF-Ca*$^{2+}$*slow/fast, BOLD-fMRI cortex/whole-brain).* ***B.*** *Standard deviation from the 'grand' averages displayed in* ***Fig. 3A.****, for each condition and group.*

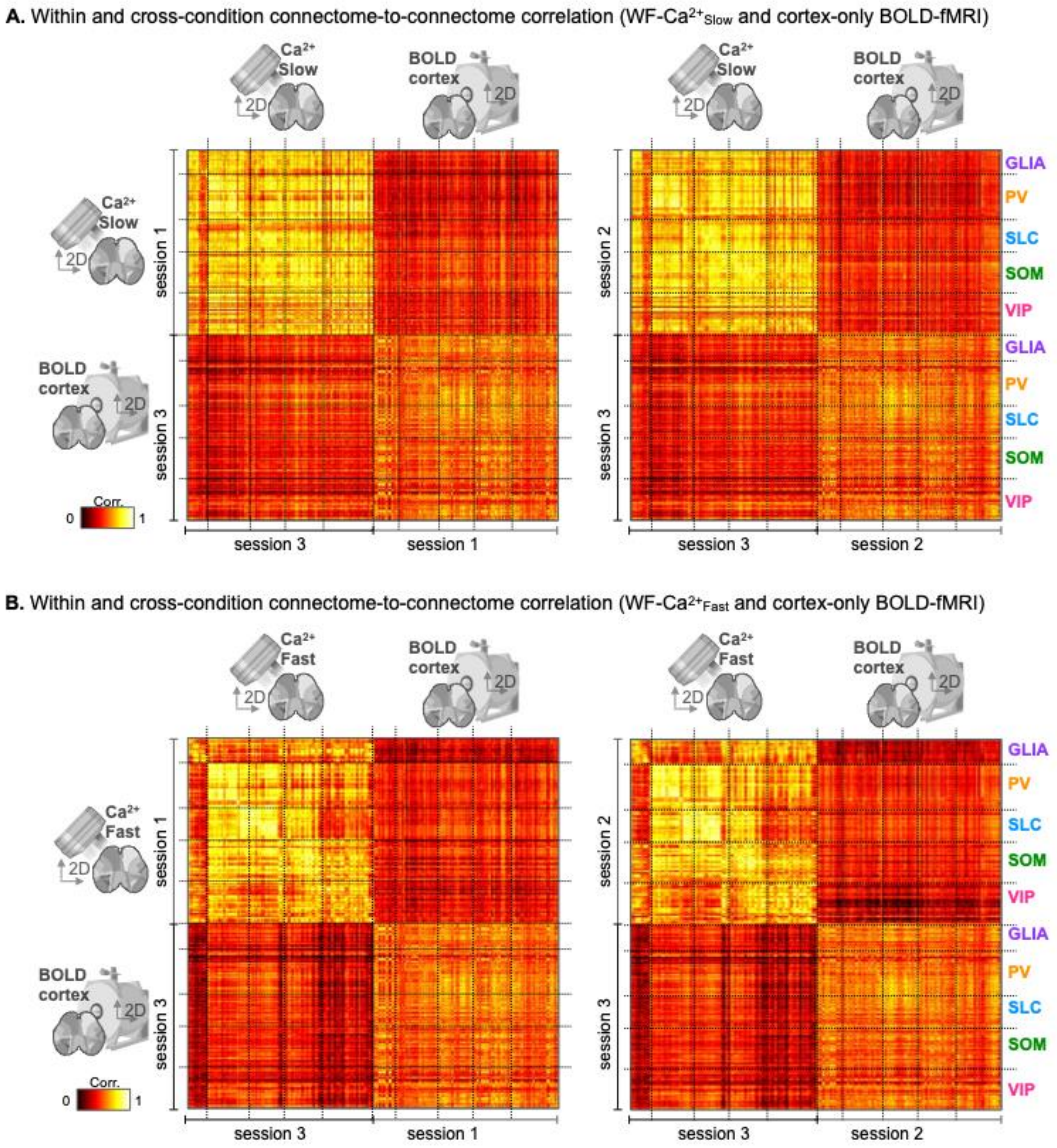


***Fig. S4* | *Within and between condition inter-connectome correlation. A.*** *WF-Ca$^{2+}$$_{Slow}$ data and BOLD-fMRI cortex-only data for session 1 ↔ 3 and 2 ↔ 3. Session 1 ↔ 2 is shown in* ***Fig. 4. B.*** *WF-Ca$^{2+}$$_{Fast}$ data and BOLD-fMRI cortex-only data for session 1 ↔ 3 and 2 ↔ 3. Likewise, session 1 ↔ 2 is shown in* ***Fig. 4.*** *Inter-connectome correlation strengths within (upper left and lower right quadrants) and between (upper right and lower left quadrants) conditions using data from within or between sessions, respectively, such that within condition comparisons were across time (i.e., from different sessions), and between condition comparisons used simultaneously collected multimodal data. Each row/column in these matrices is from a single run (i.e., a 10-minute acquisition). Inter-connectome comparisons were organized by group (top-to-bottom: GLIA, PV, SLC, SOM, & VIP) as indicated (far right).*

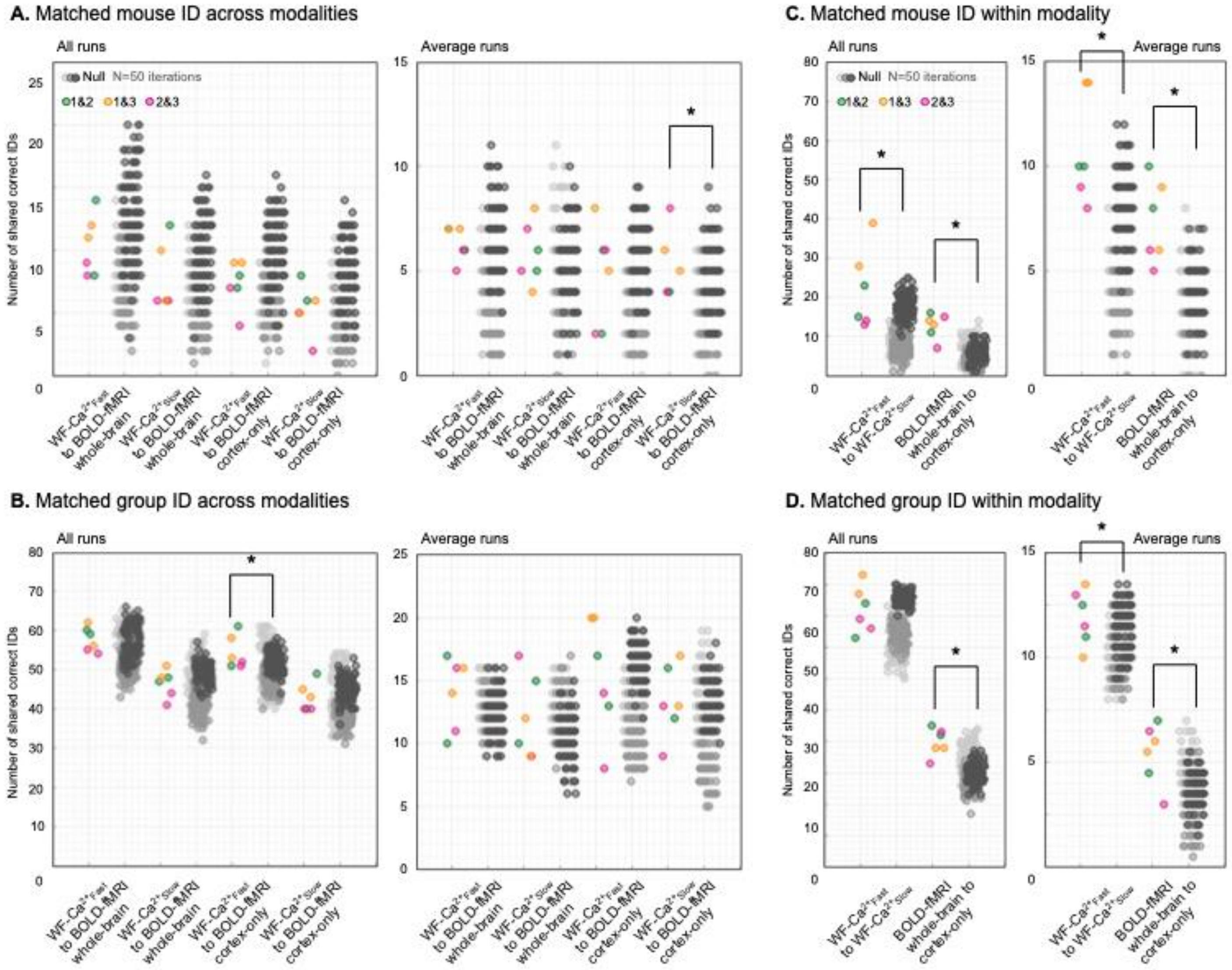


***Fig. S5 | Matching of correct mouse or group IDs across conditions between and within modalities.*** *Are the same mice (**A.** & **B.**) or group (**C.** & **D.**) IDs that are correct, using a winner-take-all framework, for one condition the same for another condition? Correct ID vectors from conditions were compared pairwise against a null generated by shuffling the correct ID vector from one of the two conditions (N = 50 iterations). Statistically significant differences (Post hoc multiple comparison (MATLAB, multcompare) results indicated with *.*

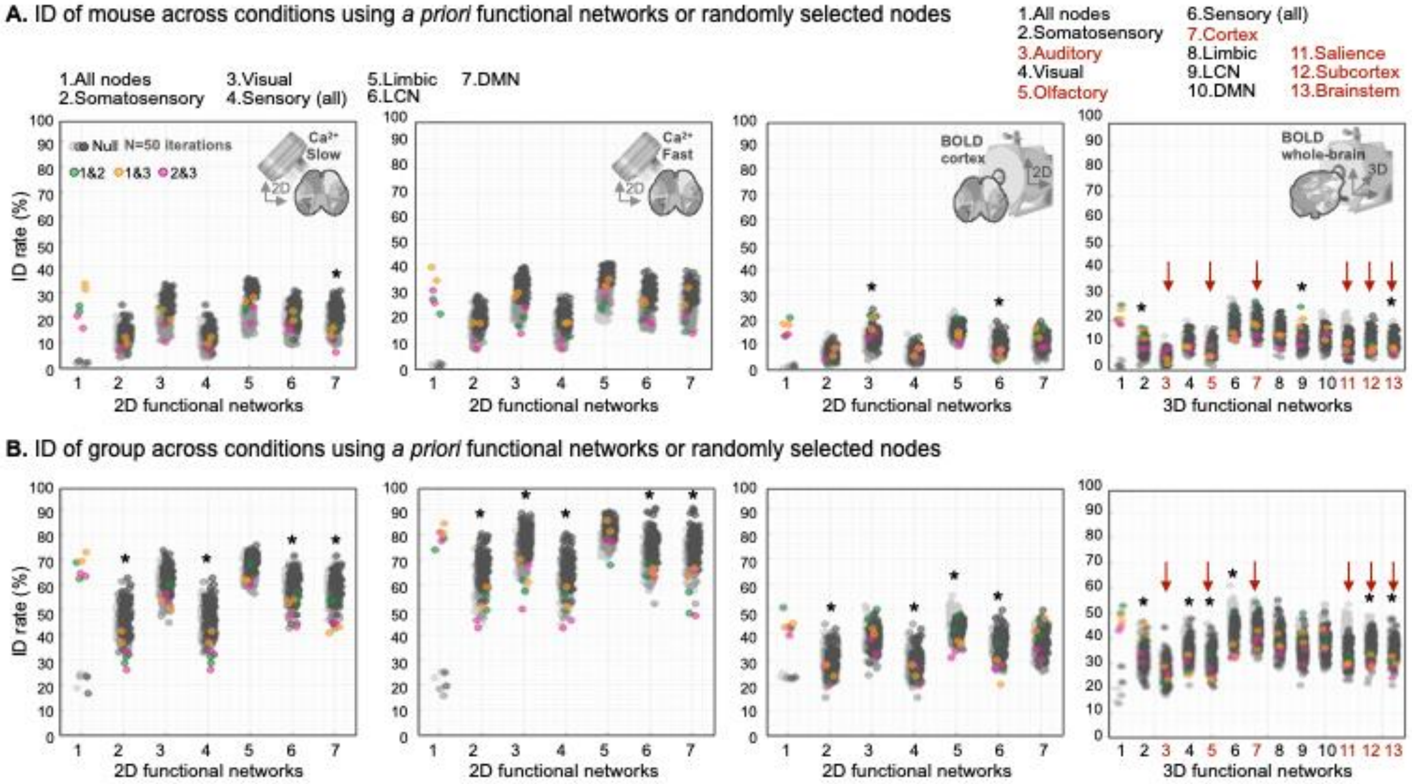


***Fig. S6* | *Intra-condition mouse and group ID using canonical resting state networks.*** *As in **Fig. 1**, mouse or group ID was repeated using randomly selected nodes ranging from 5 to all-nodes within the atlas (N = 50 iterations). In all cases, intact data outperformed shuffled data. As in **Fig. 1**, mouse **A.**, and group **B.**, IDs from random selections of nodes were compared to results obtained with a priori functional networks. See **Fig. S2** for 2D and 3D network node definitions. Thus, colored dots in **Fig. 5 A.** & **B.** (of appropriate size) are re-plotted in greyscale in **A.** and **B** here, alongside the results from canonical networks (in color). For reference, column one, in all subplots in **A.** and **B.**, were results from all nodes (color) and shuffled data (greyscale) – replicas of the rightmost columns in all subplots within **Fig. 5 A. & B.** – to bound expected high and low rates of ID for each condition. Statistically significant results in **A.** & **B.** Post hoc multiple comparison (MATLAB, multcompare with default "Tukey-Kramer test" to compute the p-values) was used after ANOVA test showed significance. Significant results indicated with *.*

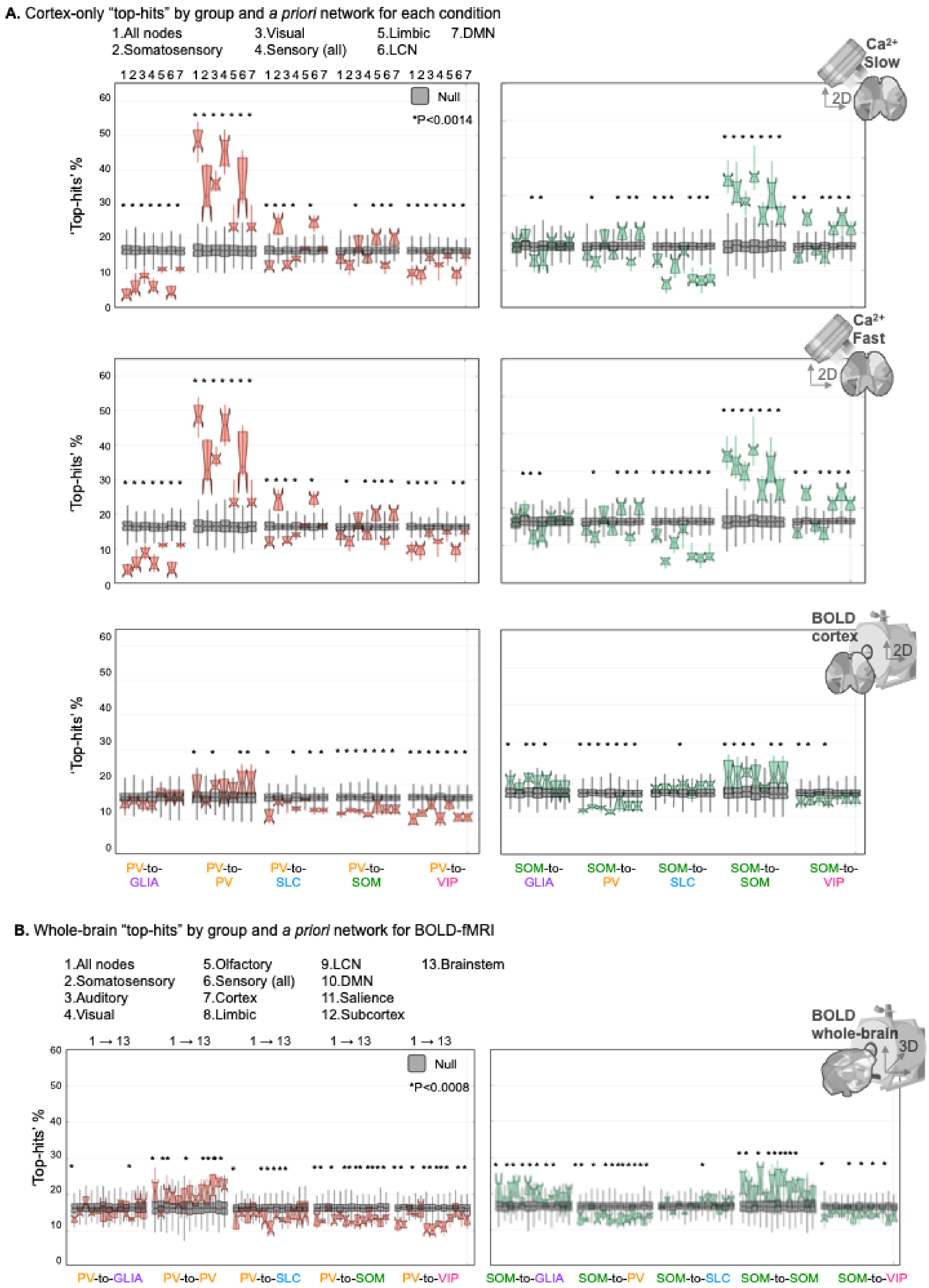


***Fig. S7 | Network influence on intra-condition ID separated by group (for PV and SOM).*** *Boxplots are constructed as described in* ***Fig. 6C.*** *Displayed as in* ***Fig. 7****. Group pairings where a statistically significant (Post hoc multiple comparison (MATLAB, multcompare) surplus, or paucity, of 'top-hits' were found are indicated by an *.*

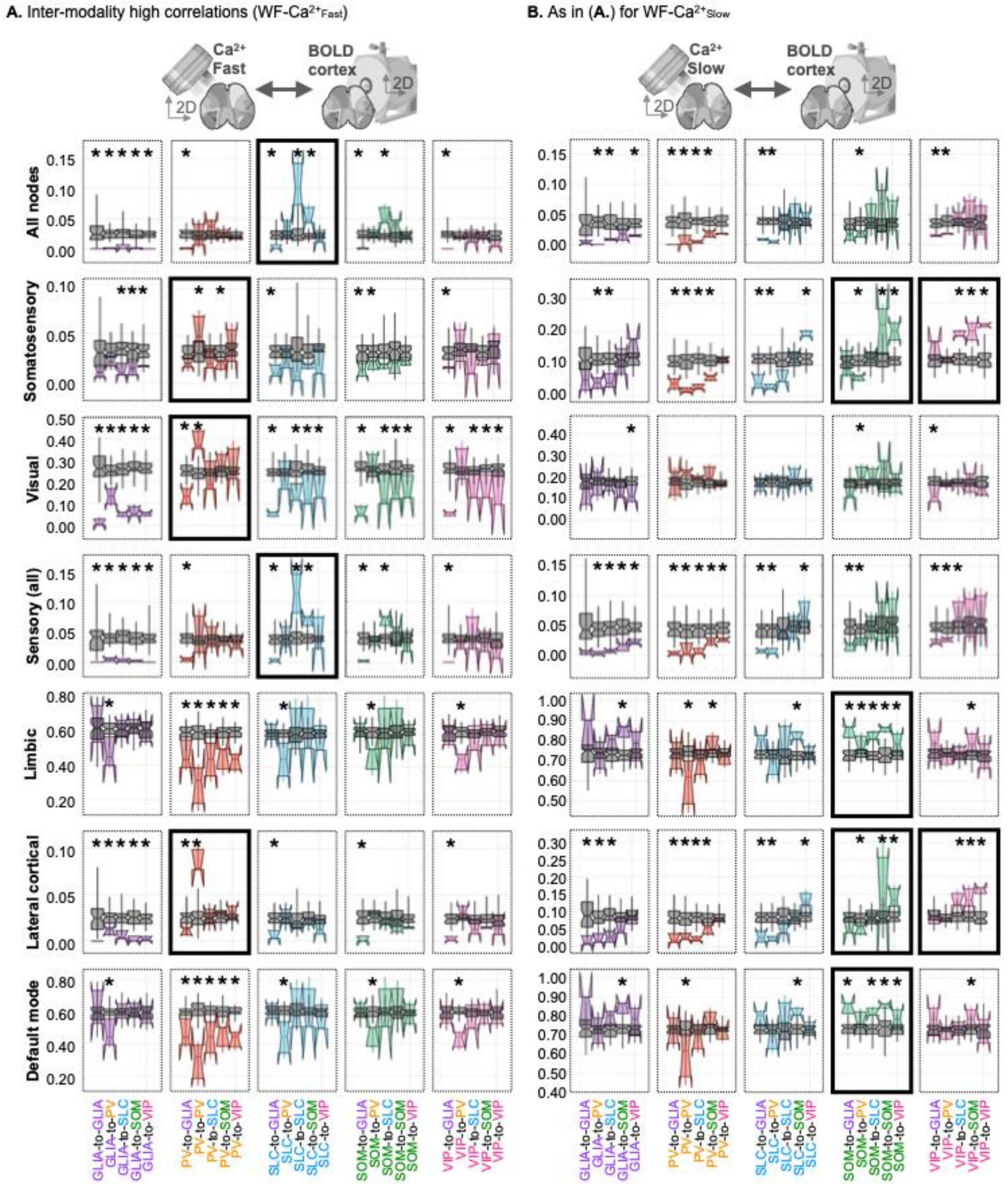


***Fig S8 | Inter-modality high connectome-to-connectome correlation strengths by network and group pairing. A.*** *Inter-modality instances of high correlation values for BOLD-fMRI cortex-only → WF-Ca$^{2+}{}_{Fast}$.* ***B.*** *Same as* ***A.****, for BOLD-fMRI cortex-only → WF-Ca$^{2+}{}_{Slow}$. Subplots where matched group pairings (e.g., GLIA-to-GLIA) showed a significant result are outlined with a solid black line. A statistically significant (Post hoc multiple comparison (MATLAB, multcompare) surplus, or paucity, of high correlation strengths are indicated by an *.*

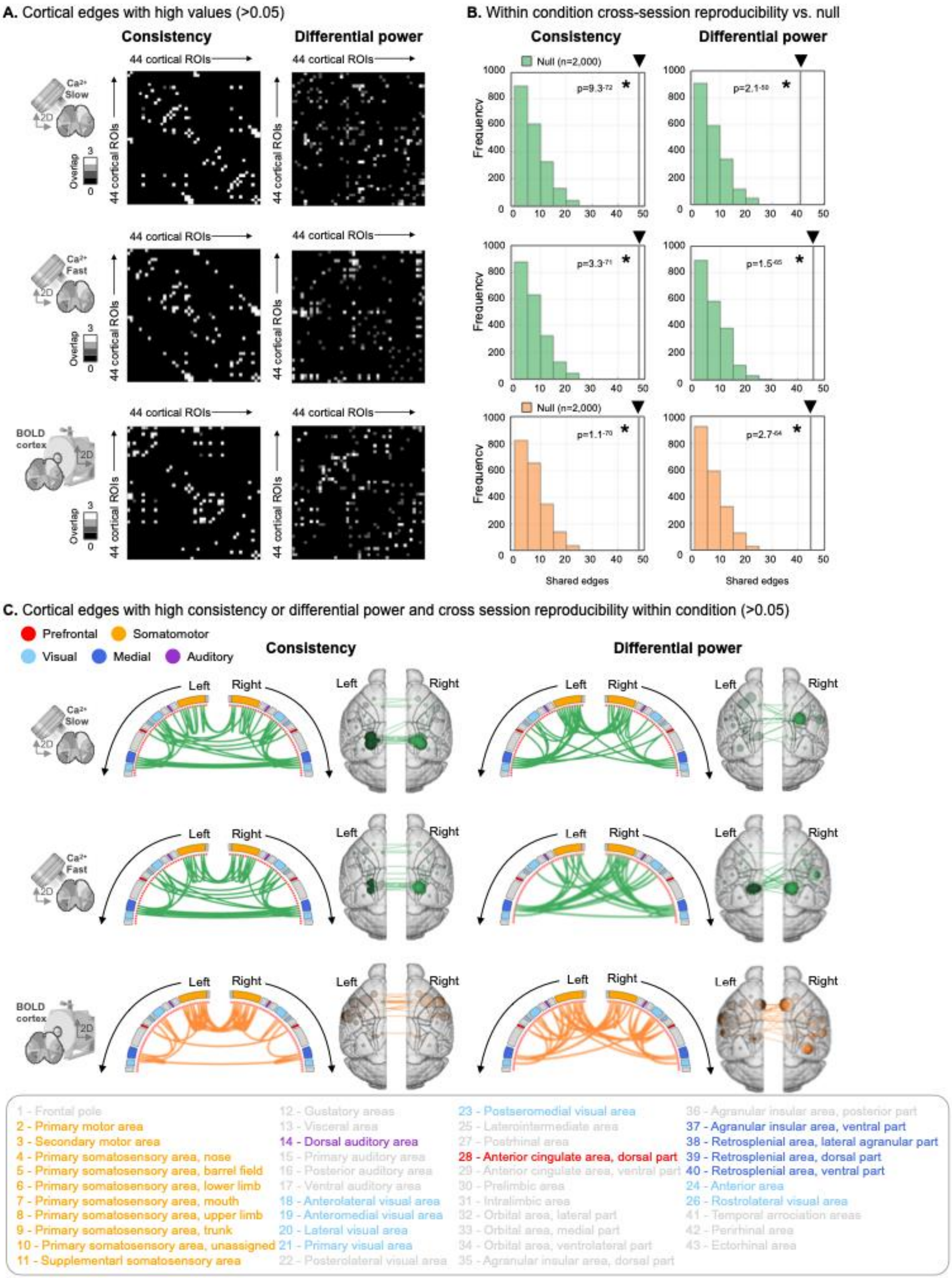


***Fig. S9. Inter-session reproducibility of high consistency or differential power edges within condition.*** *For each condition, using the 44-node cortical mask (**Fig. S2**), edges with high consistency and differential power were computed for three session pairs (sessions 1 & 2, 1 & 3, and 2 & 3), as described previously (see Bergmann et al.). For each pair of sessions, edges with high consistency (left) or differential power (right) were binarized at p<0.05 and summed across session pairs (**A.**). The number of edges with high consistency (left) or differential power (right) which appeared in more than one session pair (black line with black triangle) were compared to a null distribution (n=2,000 iterations, see **Methods**), and found to be highly overlapping (MATLAB ttest2) (**B.**). Edges with high consistency (left) or differential power (right) that were shared across session pairs displayed as circle plots or on glass brains. Regions belonging to five a priori brain modules (see Bergmann et al.) are indicated by color-coding (**C.**) (see **Fig. S10**).*

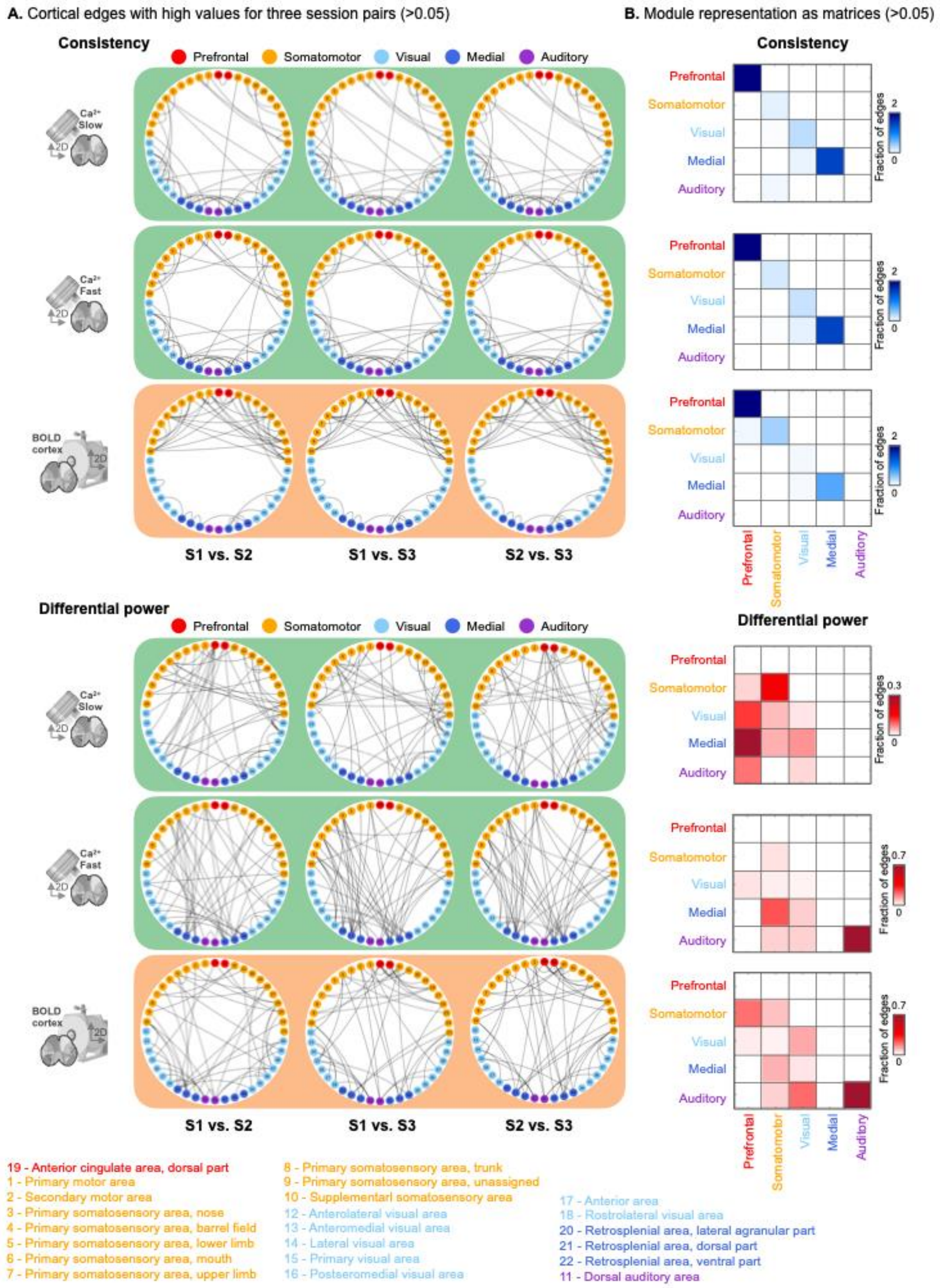


***Fig. S10. Cortical edges by brain module with high consistency or differential power within condition.*** *For each condition, using the 44-node cortical mask (**Fig. S2**), edges with high consistency (top) and differential power (bottom) are shown for three session pairs (p<0.05). Edges are displayed using circle plots and color coded using five a priori brain modules (see Bergmann et al.) (**A.**). The fraction of edges connecting between and within module for each condition are summarized in matrices (see Bergmann et al.) (**B.**).*

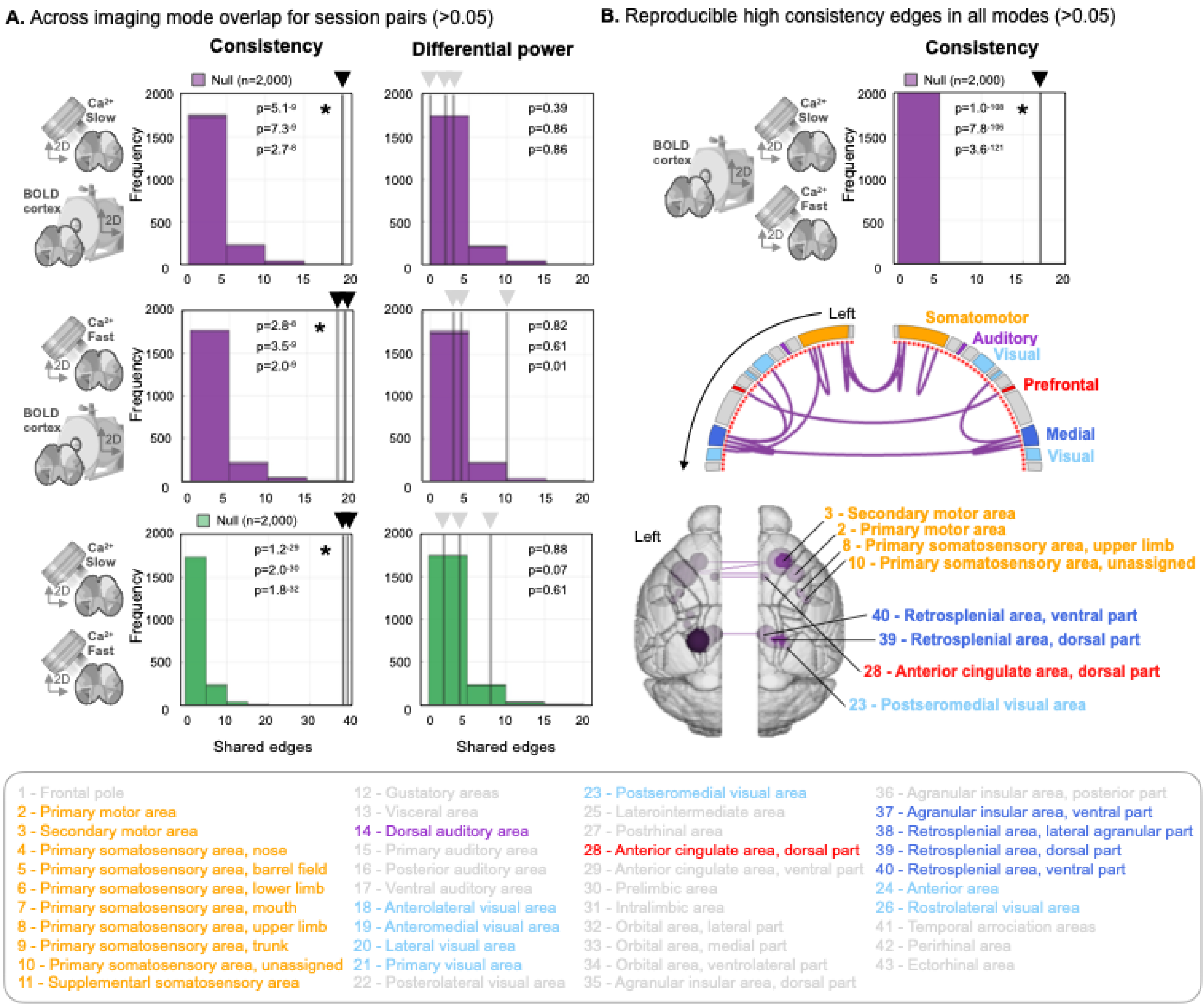


***Fig. S11. Cortical edges with high consistency or differential power shared across conditions.*** *For each pair of conditions, the number of edges with high consistency (left) or differential power (right) which appeared in both conditions (black lines with black or grey triangles) were compared to a null distribution (n=2,000 iterations, see* ***Methods****). Session pairs were considered independently and are all plotted. Edges with high consistency (left) were found to be shared across all condition pairs (and all session pairs) while edges with high differential power were not (MATLAB ttest2) (****A.****). Edges with high consistency, were shared across all three conditions (in all session pairs) (top), are displayed using a circle plot (middle), and on a glass brain (bottom). Color coding uses five a priori brain modules (see Bergmann et al.) (****B.****).*

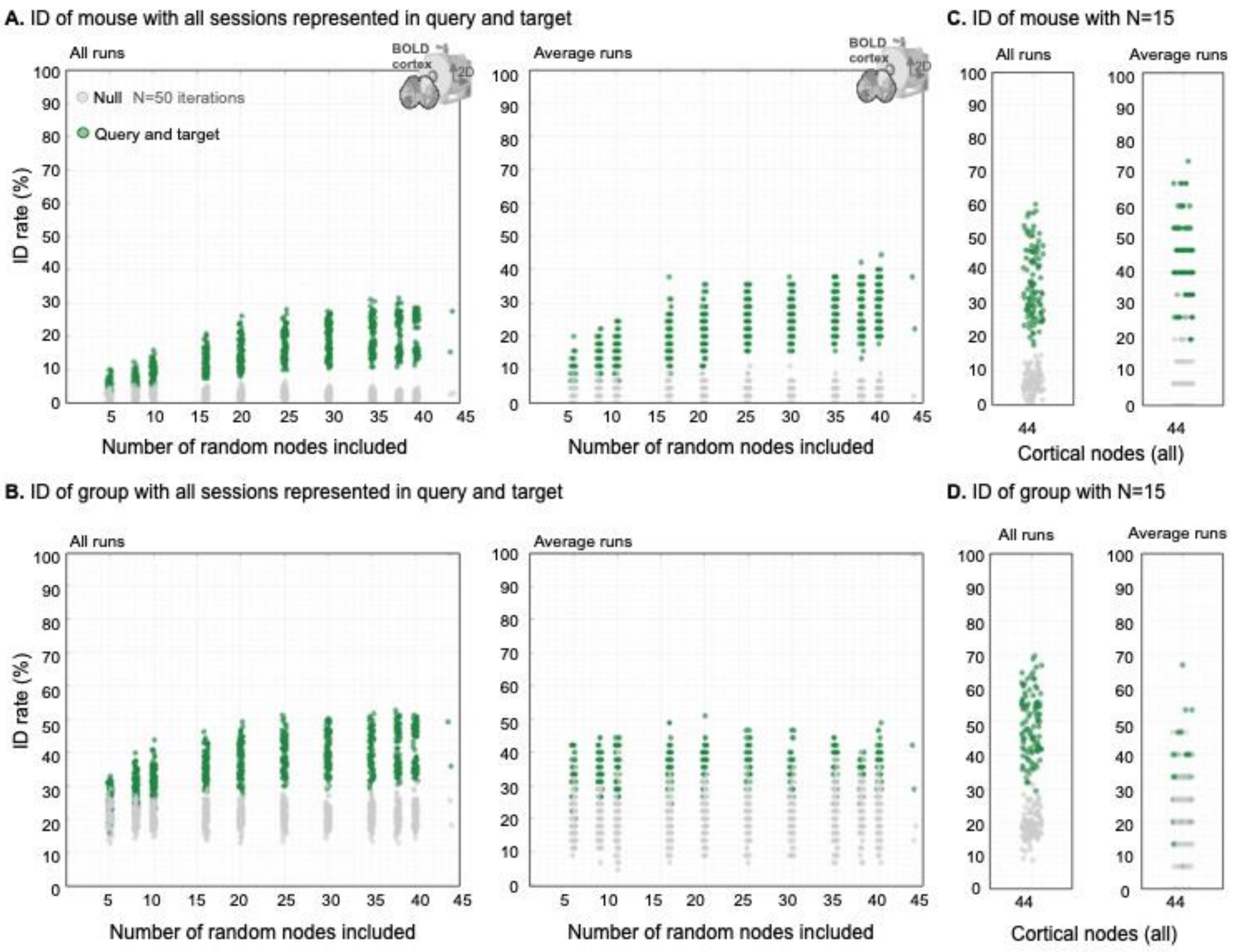


***Fig. S12 | Analysis to parallel Bergman et al. 2020.*** *Results displayed as in **Fig. 5** (**A.** & **B.**). Mouse or group ID in a winner-take-all framework with a target and query set including (non-overlapping) data representing all sessions. Similarly, (**C.** & **D.**), results from N=15 mice (balanced by group, i.e., N=3 mice per group).*

**SUPPLEMENTARY TABLES**

**Table S1.** Genotype-groups

| Offspring | Parent 1 | Parent 2 | Numbers breakdown N = animal, n = runs |
|---|---|---|---|
| **SLC** (Slc17a7-cre/Camk2α-tTA/TITL-GCaMP6f or Slc17a7-cre/Camk2α-tTA/Ai93) | Slc17a7-cre | Camk2α-tTA/TITL-GCaMP6f (Camk2α-tTA/TITL-GCaMP6f - TIGRE1.0) | N = 9, session 1/2/3 n = 26/30/30 |
| **PV** (PV-cre/Ai162) | PV-cre | Ai162 | N = 9, session 1/2/3 n = 33/29/25 |
| **VIP** (VIP-cre/Ai162) | VIP-cre | Ai162 | N = 12, session 1/2/3 n = 38/36/44 |
| **SOM** (SOM-cre/Ai162) | SOM-cre | Ai162 | N = 9, session 1/2/3 n = 29/29/32 |
| **GLIA** (Aldh1l1-creER/Ai162) | Aldh1l1-creER | Ai162 (Ai162 - TIGRE2.0) | N = 6, session 1/2/3 n = 20/20/16 |

**Table S2.** Area under curve of slow frequency content (0.008-0.2Hz). Data are shown as percentages.

| Cell type | Session 1 | Session 2 | Session 3 |
|---|---|---|---|
| **GLIA** | 77.50 | 69.40 | 58.37 |
| **PV** | 56.45 | 47.44 | 49.60 |
| **SLC** | 27.69 | 26.61 | 22.89 |
| **SOM** | 55.86 | 37.69 | 37.20 |
| **VIP** | 70.32 | 61.35 | 68.57 |

**Table S3.** Area under curve of fast frequency content (0.4-4Hz). Data are shown as percentages.

| Cell type | Session 1 | Session 2 | Session 3 |
|---|---|---|---|
| **GLIA** | 1.07 | 1.07 | 2.56 |
| **PV** | 16.72 | 19.49 | 16.98 |
| **SLC** | 34.03 | 36.28 | 43.90 |
| **SOM** | 10.49 | 16.36 | 13.51 |
| **VIP** | 4.02 | 5.51 | 5.58 |
